# Quantitative effects of co-culture on T cell motility and cancer-T cell interactions

**DOI:** 10.1101/2024.03.15.585166

**Authors:** Xinyue Li, Taoli Jin, Lisha Wang, Ming Li, Weijing Han, Xuefei Li

**Affiliations:** Key Laboratory of Quantitative Synthetic Biology, Shenzhen Institute of Synthetic Biology, Shenzhen Institutes of Advanced Technology, Chinese Academy of Sciences, Shenzhen 518055, China; Songshan Lake Materials Laboratory, Dongguan, Guangdong, 523808 China; Beijing National Laboratory for Condensed Matter Physics and CAS Key Laboratory of Soft Matter Physics, Institute of Physics, Chinese Academy of Sciences, Beijing, 100190 China

**Keywords:** T cell motility, Image data analysis, Agent-based models, Cancer-T cell interactions, Transcriptomic profiling

## Abstract

One of the primary challenges in current cancer immunotherapy is the insufficient infiltration of cytotoxic T cells into solid tumors. Despite ongoing investigations, the mechanisms restricting T cell infiltration in immune-cold tumors remains elusive, hindered by the intricate tumor microenvironment. Here, we co-cultured mouse cancer cell lines with cancer-specific cytotoxic T cells to study the influence of cancer-T cell interactions on T cell motility, a crucial factor for effective tumor infiltration. By quantifying T cell motility patterns, we found that cancer-specific T cells exhibited extended contact time with cancer-cell clusters and higher directional persistence than non-specific T cells. Computational modelling suggested that T cells with stronger persistence could facilitate efficient searching for cancer clusters. Transcriptomic profiling revealed T cells recognizing cancer cells orchestrate accumulation on cancer cell clusters by activating adhesion proteins on both cancer cells and T cells, thereby fostering prolonged interaction on cancer cells. Furthermore, we observed that there were two distinct subpopulations of cancer cells after co-culturing with cancer-specific T cells: one expressing elevated levels of T-cell attractants and antigen-presentation molecules, while the other expressing immunosuppressive molecules and undergoing epithelial-to-mesenchymal transition. These dynamic insights into the complex interplay of cancer-T cell interactions and their impact on T cell motility hold implications for refining more efficacious cancer immunotherapy strategies.

## Introduction

Cancer remains a critical global health challenge, with its devastating impact on individuals and societies. In recent years, the advent of cancer immunotherapy has revolutionized cancer treatment, offering unprecedented hope and promise for patients (1–3). Cytotoxic T lymphocytes (CTLs, CD8^+^) have emerged as a pivotal force against cancer due to their remarkable cytotoxic capabilities (4, 5). However, challenges persist in optimizing the effectiveness of immunotherapies (6). For instance, cancer cells can exploit mutations to evade immune surveillance, diminishing the efficacy of immunotherapy (7, 8). Additionally, various components such as tumor-associated macrophages (9), myeloid derive suppressing cells (10), cancer-associated fibroblasts (11), *etc.* can also impede CTLs infiltration and activation, thereby reducing patient response to immune checkpoint inhibitor therapy (12, 13). Overall, the intricate mechanisms underlying cellular interactions within the tumor microenvironment and the limitations of CTLs infiltration into solid tumors remain to be clarified.

Moreover, the spatial distribution of CTLs within the tumor microenvironment is complex and dynamic, with their motility playing a critical role in infiltration into solid tumors and anti-tumor activity. However, the impact of cancer cells on T cell motility remains poorly understood. Intrinsic factors like self-gene expression regulation (14) as well as extrinsic components such as interactions with macrophages (15) or the extracellular matrix (16), can influence T cell motility patterns. Understanding these patterns is crucial for overcoming challenges posed by the tumor microenvironment (17, 18).

However, it is technically challenging to quantitatively characterize cellular behaviors *in vivo*, particularly due to the requirement for intravital high-resolution real-time imaging techniques and the limitation of imaging period (19). Fortunately, several *in vitro* co-culture systems have been developed to investigate the interactions between T cells and tumors (20, 21), as well as T cells and extracellular matrix fibers (22, 23). These systems provide valuable tools for studying T cell mobility patterns and their interactions within controlled environments.

In this work, our objective was to investigate the movement of CTLs in the context of cancer-T cell interactions using a quasi-2D co-culture system. Through the combination of live cell microscopy, quantitative imaging data analysis, computational simulations, and transcriptomic profiling, we aimed to unravel the diverse motility patterns of cytotoxic T cells around cancer cell clusters, explore the impact of these patterns on CTL motion and spatial distributions, and gain insights into the molecular mechanisms governing these behaviours. The findings from this study not only hold significant implications for our understanding of cancer-T cell interactions on T cell motility within the tumor microenvironment, but also provide valuable data support for the advancement of immunotherapy strategies. Furthermore, these findings contribute to the broader understanding of immune response heterogeneity across different cancer cells and have implications for personalized cancer treatment approaches.

## Results

### A time-lapse pipeline for both motility and gene-expression of CTLs

The main aim of this work is to investigate the time-dependent interactions between cancer cells and cancer-specific CTLs and their impact on the mobility patterns, spatial distribution, as well as gene-expression of CTLs. To achieve this, a comprehensive study pipeline was devised, integrating a time-lapse imaging platform in a co-culture system, cell tracking and trajectory analysis, computational modeling based on cell mobility statistics, and the analysis of time-dependent gene expression in both cancer cells and CTLs (**Figure 1A**).

**Figure 1.**
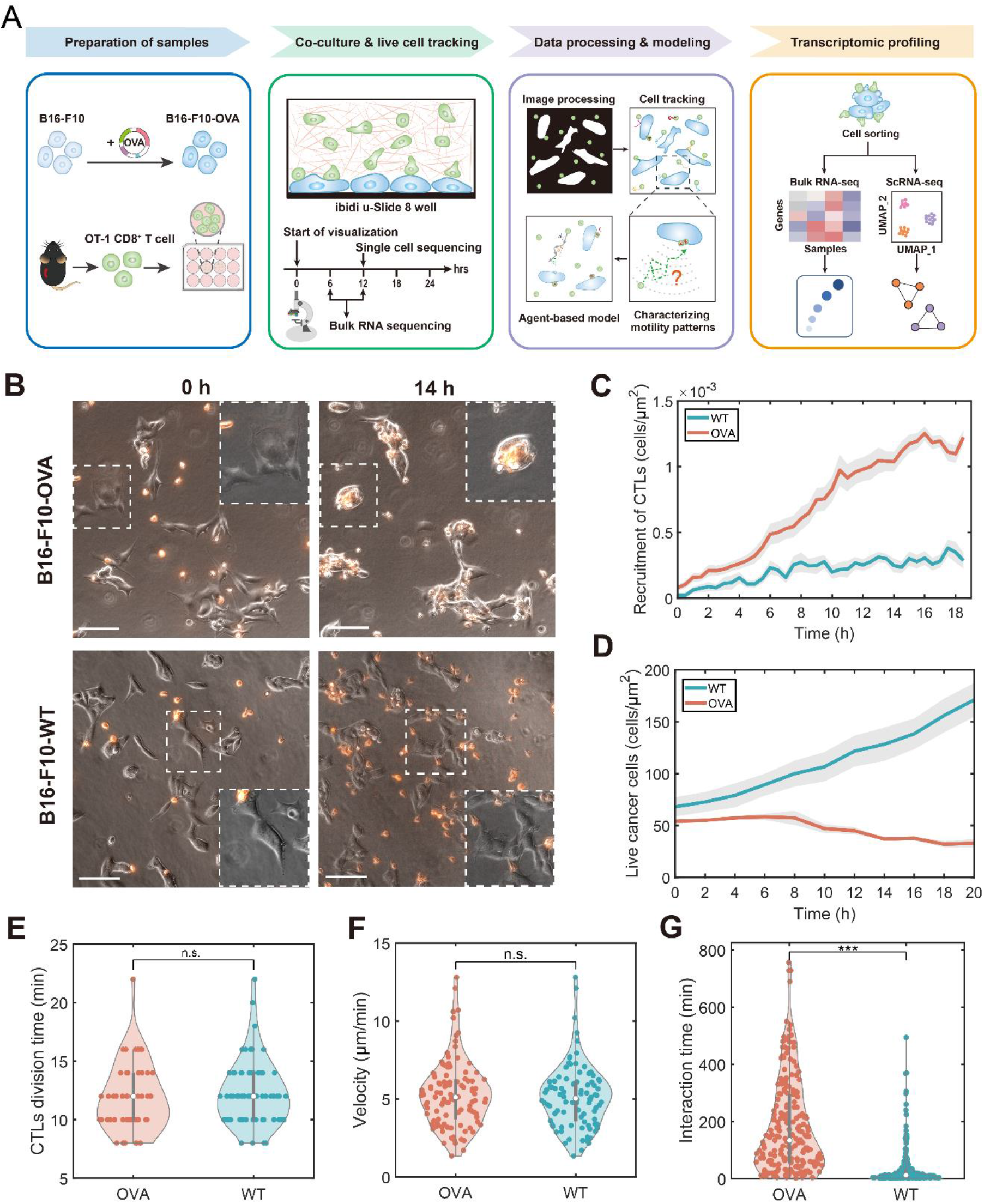
Overview of the co-culture system and pipelines. (A) Schematic representation illustrating the integrated workflow for experimental set-up, motility analysis, and gene expression profiling. *(B)* 2D microscopic images of anti-CD45-PE-labeled OT-1 CTLs (red) with murine melanoma cells (upper, B16-F10-OVA; bottom, B16-F10-WT) after coculturing 0 h and 14 h (the wide field inside the white dashed window represents an enlarged view of the small field). Scale bars, 100 μm. *(C)* Quantification of the number of recruited CTLs on the cancer cells over time for WT or OVA groups, manually determined in a 15 µm surrounding region per 30 minutes. (10∼20 regions of interest for each time point from n=1 independent experiment, mean±s.e.m.). *(D)* Counts of live cancer cells in the imaging field over time for WT or OVA groups (n=1 independent experiments, mean±s.e.m.). *(E)* Quantification of CTLs division time for B16-OVA or B16-WT groups (n = 53 vs. 69). *(F)* Migration velocity of T cells for B16-OVA or B16-WT groups (n = 107 vs. 94). *(G)* Interaction time of CTLs on cancer clusters for B16-OVA or B16-WT groups (n = 217 vs. 225). *(E-G)* Statistical significance was analyzed by unpaired t-test.

Qualitatively, under the microscope, the most apparent observation was cancer cells could be recognized by CTLs labelled by the red florescence in **Figure 1B (see also Movie S1 & Movie S2)**. To quantitatively demonstrate such accumulation, the density of T cells within the regions of interests was manually counted every 30 minutes after the establishment of co-culture (see **Methods**). Not surprisingly, the density of CTLs in the co-culture with OVA cancer cells increased with time at a rate faster than that of CTLs co-cultured with WT cancer cells (**Figure 1C and Dataset S1**). In addition, the WT cancer cells demonstrated continued growth in the co-culture with CTLs, whereas the number of OVA cancer cells gradually decreased (**Figure 1D and Dataset S1**).

To elucidate the faster accumulation of CTLs shown in **Figure 1C**, we first hypothesized that this could be due to the faster growth or swifter mobility of CTLs co-culture with OVA cancer cells. However, no significant differences were observed in the doubling time or velocity between CTLs in the two co-cultures (**Figure 1E**, **Figure 1F and Dataset S1**). Alternatively, such accumulation of CTLs could also be attributed to their reduced mobility around the OVA cancer cells or a more pronounced chemotactic motion towards OVA cancer cells. Intriguingly, manually tracking CTLs on both types of cancer cells revealed an extended engagement between CTLs and OVA cancer cells (**Figure 1G and Dataset S1**). Due to the constraints of manually handling CTL tracks, in the following analysis, we turned to an automated tracking strategy to investigate the potential chemotaxis behavior of CTLs.

### Quantitative motility patterns of antigen-specific T cells

To quantitatively investigate the motility behaviours of CTLs in the co-culture with OVA and WT cancer cells, our focus was on characterizing their motility patterns before and after CTLs reached cancer cell clusters. The procedure for time-lapse image processing and data analysis is illustrated in **Figure 2A** and **2B** (see **Methods** and **Fig. S1**). Utilizing the automated time-lapse image data processing pipeline, a total of 4276 and 5146 CTL tracks were collected from 5 parallels in the OVA and WT groups, respectively, throughout the experimental observation period. Subsequent analyses were conducted based on the identified tracks of CTLs and the corresponding areas of cancer-cell clusters.

**Figure 2.**
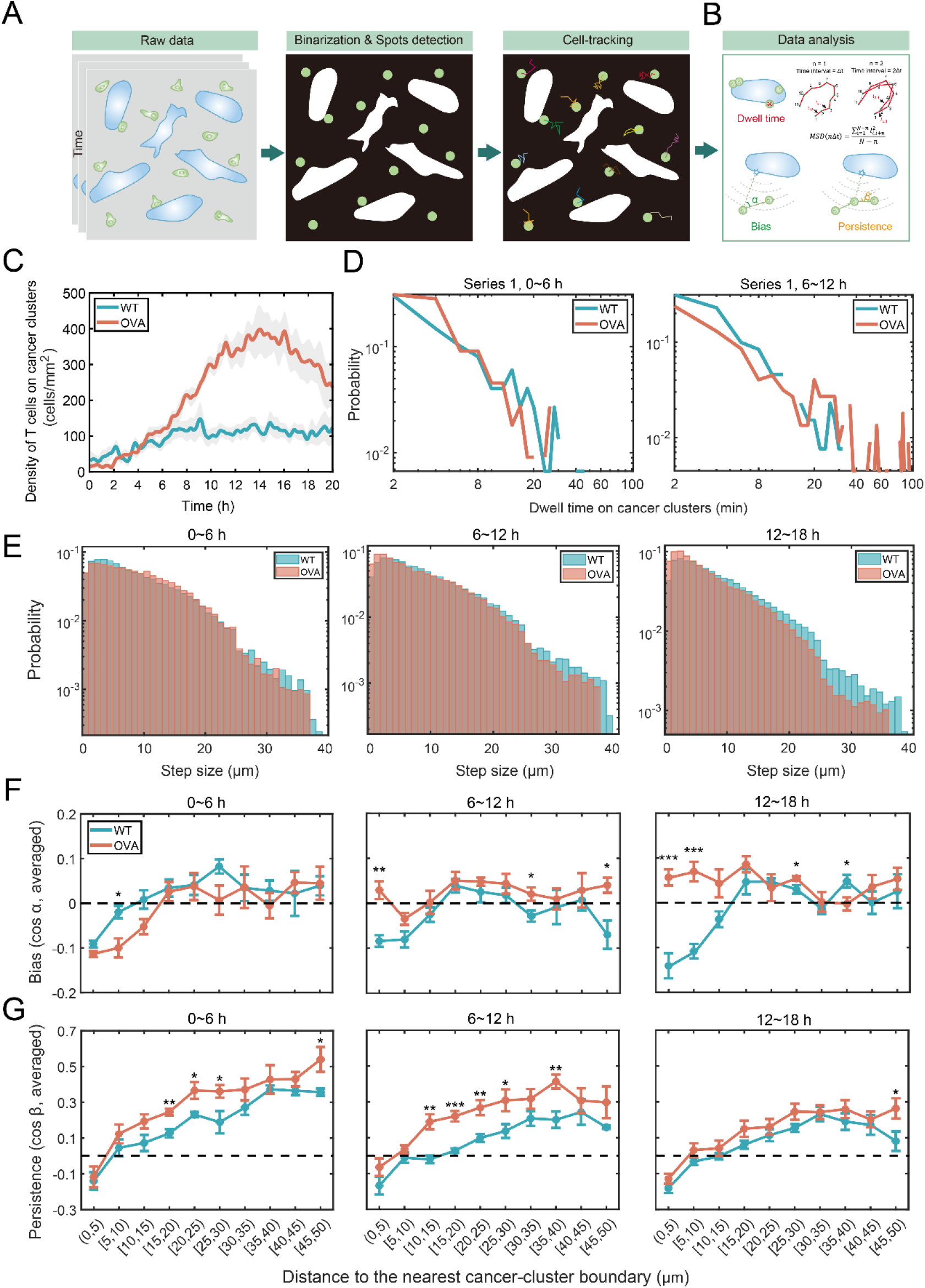
Quantitative motility patterns of CTLs in the co-culture with OVA and WT cancer cells. *(A)* Pipeline for processing time-lapse images and tracking T cells. *(B)* Schematic representation of data analysis, including dwell time, mean square displacement, directional bias, directional persistence of CTLs towards cancer cell clusters. *(C)* Density of CTLs on cancer clusters (cells/mm^2^) over time, demonstrating a significant T cell accumulation in OVA groups compared to WT groups. *(D)* Dwell time distribution of T cells on cancer-cell clusters in Series 1 within 6-hour intervals. Left, 0-6 h; right, 6-12 h. The distribution for 12-18 h is shown in **Fig. S4.** *(E)* Step sizes distributions of CTLs within 6-hour intervals. Left, 0-6 h; middle, 6-12 h; right, 12-18 h. *D-E*, the x and y axes were scaled logarithmically. *(F)* Directional bias or *(G)* persistence distributions of CTLs movement towards the nearest cancer cell boundary within 3 time-intervals. Left, 0-6 h; middle, 6-12 h; right, 12-18 h. *p*<0.001 (***), *p*<0.01 (**), *p*<0.05 (*). *(C-G)*, Red lines (or bars) represent OVA groups, and blue lines (or bars) represent WT groups. *(C, F&G)*, n=5 independent experiments, mean±s.e.m.

Firstly, we quantified the behaviours of CTLs when their fluorescent signals overlapped with those of cancer cells, indicating direct contact and interactions between CTLs and cancer cells (see **Methods**). A substantial increase in the number of CTLs on OVA cancer-cell clusters was demonstrated within 6-12 h after co-culturing (**Figure 2C, Fig. S2 and Dataset S2**), which was not observed in the co-culture with WT cancer cells. This result was consistent with the manually characterized CTL-density (**Figure 1C**). In addition, the total number of CTLs per unit area over time in the imaging field for both co-culture systems showed no significant difference (**Fig. S3**), suggesting that the stronger accumulation of CTLs on OVA cancer-cell clusters might not result from differential CTL proliferation. Furthermore, to evaluate the interaction time between CTLs and cancer cells, the dwell time distribution of CTLs on cancer-cell clusters was quantified (**Figure 2D, see Methods**). The result revealed that the dwell time distribution closely followed a heavy-tailed power-law distribution. Notably, within the first 6 hours, there was no significant difference in the dwell time distributions of CTLs between the co-culture systems with OVA and WT cancer cells. However, during 6-18 hours of the co-culture, the distribution on OVA cancer-cell clusters exhibited a longer tail than on the WT cancer-cell clusters (**Figure 1D, Fig. S4, Fig. S1D and Dataset S2**), where some CTLs could stay on OVA cancer-cell clusters for up to 100 minutes. This finding indicates that once CTLs recognized and interacted with cancer cells, they tended to stay on cancer cells for a longer period.

Secondly, to investigate the motility behaviours of CTLs before they reached cancer-cell clusters, we studied the step sizes, mean square displacement (MSD), directional bias and directional persistence of CTLs when their fluorescent signals did not overlap with those of cancer cells (see **Methods**). We observed a slight difference in CTLs step size distributions as a function of distance from the cancer boundary (**Figure 2E, Fig. S6 and Dataset S2**). CTLs within 5 microns from cancer boundary exhibited smaller step sizes, typically less than 3 microns (**Figure 2E & Fig. S6**). After 12 h, CTLs co-cultured with OVA cancer-clusters showed shorter step sizes compared to those with WT cancer-clusters (**Figure 2E, right panel & Fig. S6, bottom two rows**). Moreover, CTLs demonstrated super-diffusive motion behaviours, characterized by MSD∼τ^α^, with the exponent α around 1.3 (**Fig. S7**). Intriguingly, during the first 6 h of the co-culture, no significant difference in MSD power exponents was observed between the two systems (**Fig. S7**, left panel). However, in the subsequent 6-12 h (**Figure 2D**, right panel) and 12-18 h (**Fig. S7**), CTLs co-cultured with OVA cancer cells exhibited a higher power exponent compared to those co-cultured with WT cancer cells, suggesting that CTLs co-cultured with OVA cancer cells may possess a more efficient searching capability (24).

To further investigate the motility patterns of CTLs with respect to their distance towards cancer cell clusters, directional bias and directional persistence were examined (see **Methods**, **Ref.** (**25**)). CTLs co-cultured with OVA cancer cells gradually increased their motion bias towards the nearest cancer-cell clusters from the first 6 hours to the 9-18 hours of the co-culture, particularly when 0-10 μm away from cancer-cell clusters (**Figure 2F**, **Fig. S8 and Dataset S2**). By contrast, CTLs co-cultured with WT cancer cells showed a gradual decrease in directional bias, especially within 0-10 μm of cancer-cell clusters. Therefore, although CTLs co-cultured with OVA cancer cells only displayed a mild bias (∼0.1) toward the nearby cancer clusters within 9-18 h (**Figure 2F** and **Fig. S8**), there was a significant difference compared to CTLs with OVA cancer cells, suggesting that the interactions between cancer and CTLs may play a crucial role in attracting CTLs towards the nearby cancer clusters.

For the directional persistence, CTLs co-cultured with OVA cancer cells displayed considerably stronger persistence compared to those cultured with WT cancer cells, particularly during 3-12 h (**Figure 2G, Fig. S9 and Dataset S2**). These observations indicate that the interactions between OVA cancer cells and CTLs might enhance the ability of CTLs to maintain the consistent movement direction. It is noteworthy that a consistent decrease in directional persistence over time was observed in both co-culture systems. To explain this decrease, we hypothesized that the orientation or density of collagen fibers around cancer cell clusters might affect the directional persistence. While no significant difference in the orientation of fibers around cancer cell clusters in the two co-culture systems was observed (**Fig. S10**), the density of collagen fibers consistently decreased as a function of the distance towards cancer cell clusters (**Fig. S11 and Movie S3-S6**). Therefore, we argue that as CTLs approached cancer cells, they encountered higher collagen density, leading a higher probability to change their orientation by following the fibers or penetrating through the fiber gaps, as suggested by others (26).

Furthermore, we conducted pairwise correlation tests among step size, directional persistence, and directional bias at each distance to the cancer boundary within 6-hour intervals (**Fig. S12**). No significant correlation was found between step size and directional bias. While correlations between directional persistence and bias were observed within a 10 μm distance from cancer cell clusters, they were deemed negligible. Interestingly, significant correlations between step size and directional persistence were identified beyond the 10 μm range. However, these correlations were generally weak, particularly when considering CTLs co-cultured with WT cancer cells in comparison to those with OVA cancer cells.

### Computational studies revealed the key motility patterns governing the efficient searching and the accumulation of CTLs on cancer-cell clusters

In **Figure 2**, we provided a quantitative and systematic characterization of the 2-dimentional motility patterns of CTLs. To further elucidate the roles of different T-cell motility patterns in their searching and accumulation on cancer-cell clusters, we developed a 2-dimensional agent-based model where four motility characteristics of T cells, i.e., dwell time, directional bias, persistence, and step-size distribution, were varied and their effects on the accumulation of T cells were compared (**Figure 3A**, see **Methods**), with a display of simulated trajectories in one simulation in **Figure 3B and Movie S7**.

**Figure 3.**
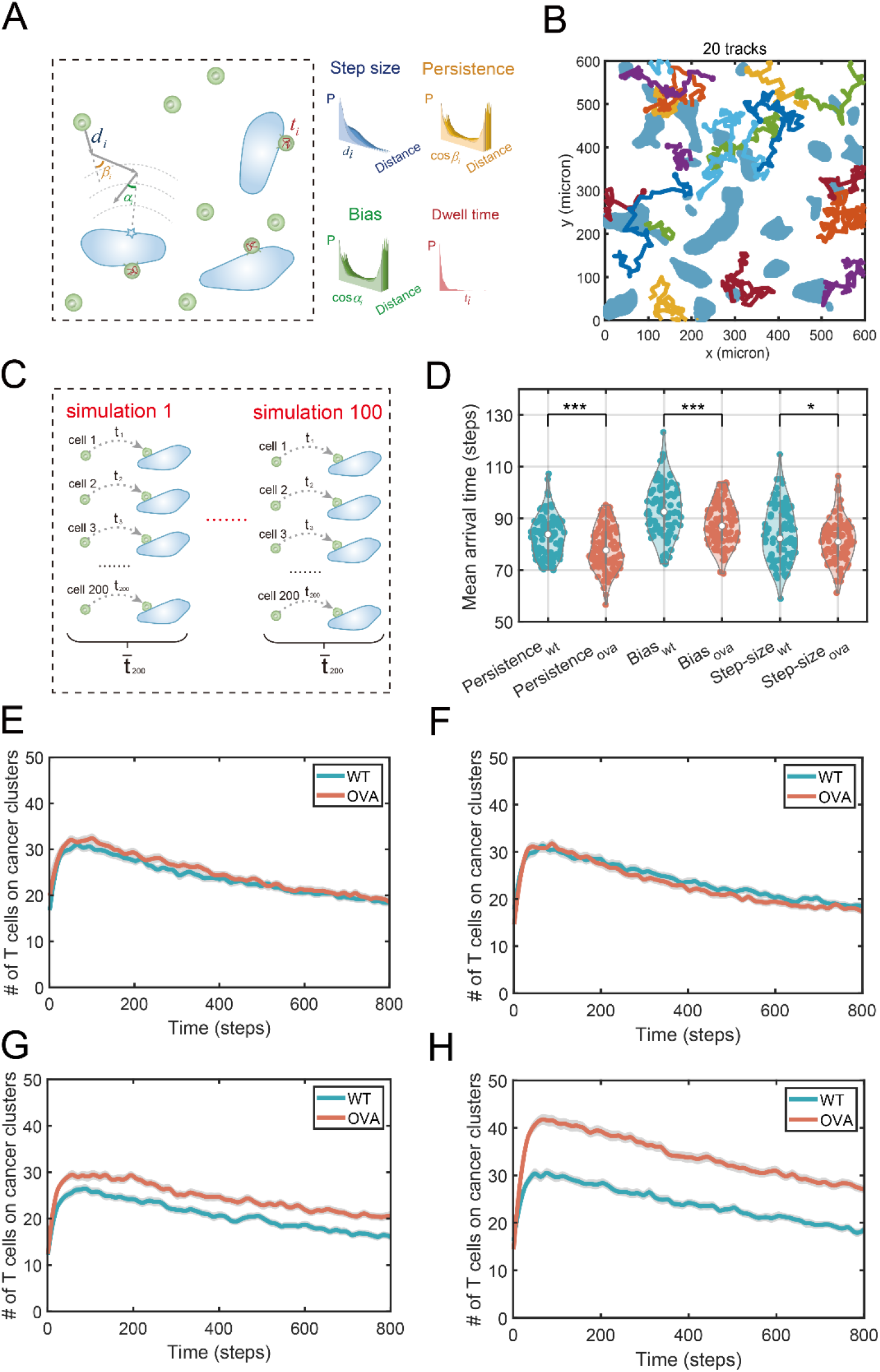
Computational studies revealed the key motility patterns for the efficient searching and the accumulation of CTLs on cancer-cell clusters. *(A)* Schematic representation of agent-based models. *(B)* An illustration depicting the trajectories of 20 CTLs after moving 100 steps each. *(C)* Schematics of the calculation method of mean arrival time of CTLs towards cancer clusters. *(D)* Violin plot and statistical significance test of the mean arrival time of CTLs to cancer clusters under different patterns when altering only the differences in the directional persistence or directional bias or step sizes. 100 replicated simulations were performed, with each simulation consisting of 200 cells moving for 800 steps. ‘*’ represents *p*<0.05, ‘**’represents *p*<0.01 and ‘***’ represents *p*<0.001. *(E-H)* The accumulation of CTLs on cancer-cell clusters as a function of time when considering different variations: *(E)* step size, *(F)* directional persistence, *(G)* directional bias, and *(H)* dwell time. The shaded area represents the 95% confidence interval of the simulated data, while the bold line represents the mean.

Firstly, we explored the impact of motility patterns on the mean arrival time of CTLs towards cancer-cell clusters (**Figure 3C**). Specifically, the distance-dependent directional persistence, bias, and step-size distribution of CTLs characterized from the two coculture systems were utilized (**see Methods**). By manipulating one pattern while maintaining consistency in the other two patterns with the co-culture system involving WT cancer cells and CTLs, we investigated whether observed differences in CTL motility patterns between the two co-culture systems could lead to variations in the mean arrival time of CTLs. Notably, statistically significant differences were observed when altering the directional persistence or bias, rather than the step-size (**Figure 3D and Dataset S3**). Intriguingly, CTLs in the co-culture with OVA cancer cells reached cancer-cell clusters more rapidly, emphasizing the importance of directional persistence and bias in CTLs’ search strategy.

In addition, to ensure the robustness of our findings, we conducted similar analyses under the condition where there was only a single cancer-cell cluster in the simulation box (**Fig. S13A-B & Movie S8**). The observed trends remained consistent when considering only the differences in directional persistence or step-size, while showing inconsistency in directional bias, which can be attributed to the initial distribution of CTLs (see **Fig. S13C**). These results suggest that stronger directional persistence could enhance CTLs’ efficiency in searching for cancer-cell clusters.

Subsequently, we investigated the impact of motility patterns on the accumulation of CTLs on cancer-cell clusters. No significant difference in the number of CTLs on cancer-cell clusters was found when varying the step-size (**Figure 3E and Dataset S3**) or the directional persistence distribution (**Figure 3F and Dataset S3**), while maintaining consistency in the dwell-time distribution with the coculture system involving WT cancer cells and CTLs. However, we did observe a slight difference in accumulation of CTLs when considering the differences in the directional bias (**Figure 3G and Dataset S3**). In contrast, a significant accumulation of CTLs on cancer clusters was observed when considering differences in the dwell-time distribution in simulations with OVA cancer cells (**Figure 3H**). Furthermore, we demonstrated that such accumulation was not qualitatively affected by the configuration of the cancer-cell clusters (**Fig. S13D-G**). These findings suggest that stronger adhesion, rather than the motion outside of cancer-cell clusters, plays a more crucial role in the accumulation of CTLs on cancer clusters.

### Potential mechanisms underlying the mobility patterns elucidated via bulk RNA sequencing analysis

To investigate the potential molecular mechanisms underlying T cell motility patterns, we performed both bulk and single-cell RNA sequencing of cancer and T cells sorted from the co-culture systems (see **Methods**).

First, by analyzing bulk RNA-seq data of T cells co-cultured with OVA cancer cells, we found that in addition to classical cytokines/factors related to T-cell activation/cytotoxicity, such as interleukin-2, perforin, interferons and tumor-necrosis factors, *etc.*, chemokines that could be responsible for recruiting/polarizing innate immune cells were also significantly up-regulated, such as Xcl1, CCL1, CSF2 (**Figure 4A**), gradually switching on after the co-culture (**Figure 4B**). Overall, 262 genes were up regulated at 12 hours after co-culture (see **Methods**). By performing the KEGG pathway enrichment analysis (**Figure 4C**) as well as the gene ontology (GO) enrichment analysis (**Figure 4D**) of these genes (see **Methods**), we found that cytokine-cytokine receptor interaction and chemokine signaling pathways were significantly enriched. Interestingly, GO analysis further revealed that up-regulated cytokines/chemokines were enriched in the recruitment of innate immune compartment, which could be essential for antigen-specific CTLs to coordinate the immune microenvironment around cancer cells.

**Figure 4.**
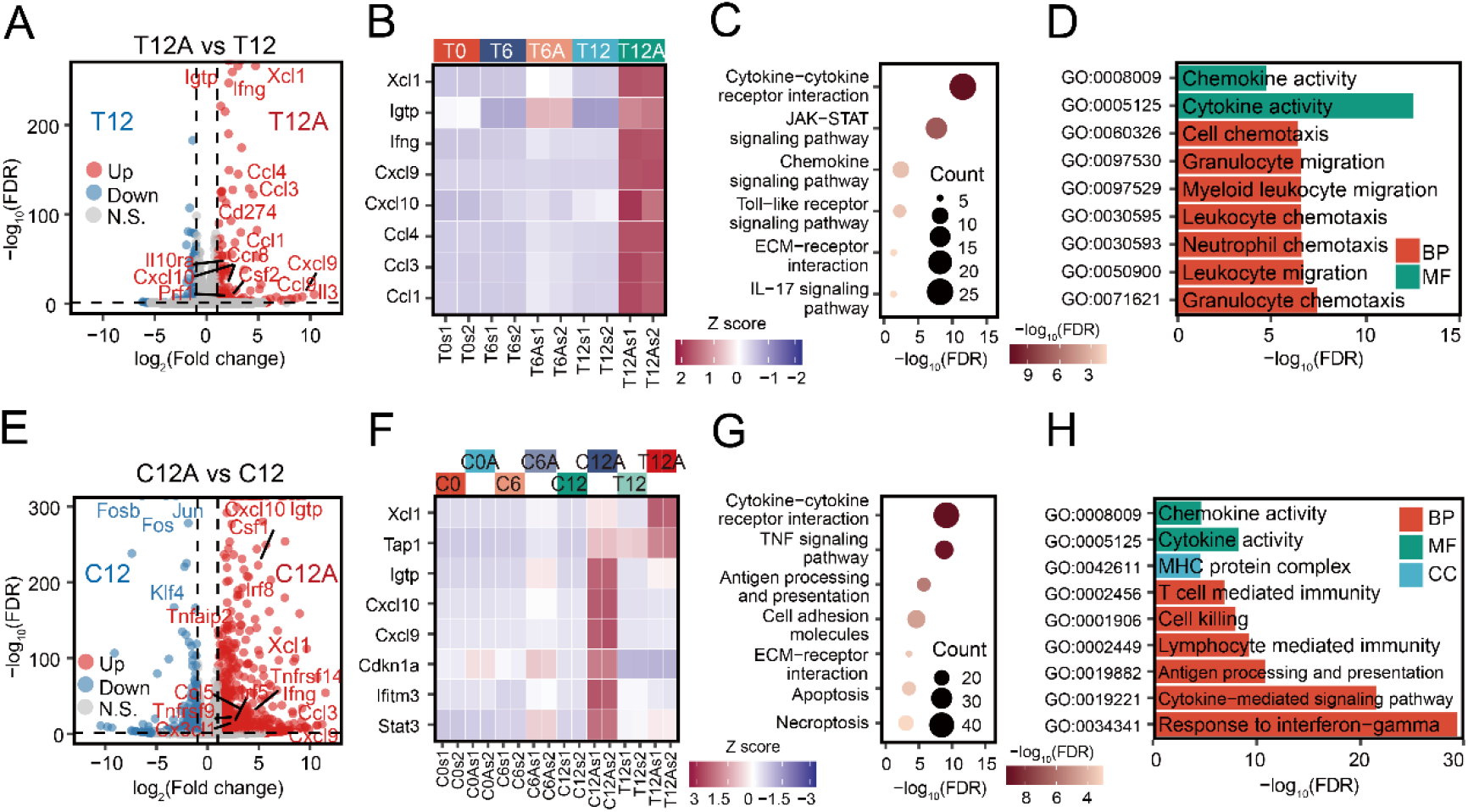
Analysis of the bulk RNA sequencing data of cancer cells and CTLs. *(A)* Volcano plot of differentially expressed genes between CTLs co-cultured with OVA cancer cells and those co-cultured with WT cancer cells at 12 h after the co-culture (T12A vs T12). Genes that were significantly up and down regulated in T12A samples, i.e., log2(fold change)>1 and FDR<0.05, were shown in red and blue, respectively. *(B)* Temporal dynamics of selected up-regulated genes in panel *A*. CTLs at 0, 6, 12 h of the co-culture with OVA or WT cancer cells were filtered and sequenced in two replicates. The Z-score of the corresponding expression of the selected genes were calculated and demonstrated. *(C)* Results of the KEGG pathway enrichment analysis of differentially expressed genes shown in panel *A*, where the size of the circle corresponds to the number genes in the specific pathway and the color corresponds to FDR values of the enrichment analysis. *(D)* Results of the gene ontology (GO) enrichment analysis of differentially expressed genes shown in panel *A*, where different colors correspond to GO terms for biological process (BP), molecular function (MF) or cellular component (CC). *(E)* Volcano plot of differentially expressed genes between OVA and WT cancer cells at 12 h after the co-culture with CTLs (C12A vs C12). Genes that were significantly up and down regulated in C12A samples were shown in red and blue, respectively. *(F)* Temporal dynamics of selected up-regulated genes in panel *E*. OVA and WT cancer cells at 0, 6, 12 h of the co-culture with CTLs were filtered and sequenced in two replicates. The Z-score of the corresponding expression of the selected genes were calculated and demonstrated. *(G)* Results of the KEGG pathway enrichment analysis of differentially expressed genes shown in panel *E*. *(H)* Results of the gene ontology (GO) enrichment analysis of differentially expressed genes shown in panel *E*.

For the gene expression profiles of cancer cells, compared to WT ones, 942 genes were up regulated in OVA cancer cells at 12 h after the co-culture with CTLs (**Figure 4E**). For specific genes, given the up-regulation of interferon-gamma by antigen-specific CTLs, it is not surprising that OVA cancer-cell genes that respond to interferon protein family were significantly up regulated, such as Stat3, Ifitm3, Cdkn1a, *etc*. Interestingly, two key chemokines for attracting T cells, CXCL9 and CXCL10, were gradually up regulated in OVA cancer cells after interactions with CTLs. It is worth noting that their expression level in OVA cancer cells is much higher than that in T cells (**Figure 4F**).

By further performing the KEGG pathway enrichment analysis (**Figure 4G**) as well as the GO enrichment analysis (**Figure 4H**) of the 942 up-regulated genes in OVA cancer cells, we demonstrated that cancer cells respond to CTL-attack by enhancing genes that are enriched in pathways such as antigen processing and presentation, apoptosis, TNF signaling pathway, *etc.*, as well as enriched in GO such as MHC protein complex, response to interferon-gamma, cell killing, *etc*. Furthermore, we noted that pathways of cell adhesion molecules and ECM-receptor interaction are enriched for genes up-regulated in OVA cancer cells (**Fig. S14A**). When looking into the specific genes, we found that some characteristic genes that were related to epithelial-to-mesenchymal transition were up regulated, such as Serpine1, Col1a1, Dab2, *etc.* After confirming the homologs corresponding to the human genes in 50 hallmark genes sets from the MSigDB, we demonstrated increased epithelial-to-mesenchymal transition (EMT, NES:1.77, FDR:5.6e-05) with the GSEA analysis (**Fig. S14B**). Consistently, GO term related to epithelial-to-mesenchymal transition were also enriched (**Dataset S4**). These observations suggest that in response to T-cell attack, cancer cells may tend to metastasize.

Furthermore, focusing on the temporal dynamics of gene expression, among the up-regulated 1239 genes (C12A vs. C0A), 137 of them (Set 1) were progressively up-regulated from 0 to 12 hours, whereas 604 genes (Set 2) were promptly boosted at 6 hours and kept relatively stable until 12 hours (**Dataset S4**). Inspired by such differential temporal dynamics of the gene expression, we further tested which genes in Set 2 have a regulatory role on genes in Set 1 (see **Methods**). And we found that 8 genes encoding corresponding transcription factors in Set 2 can enhance the expression of 13 genes in Set 1 (**Dataset S4**). The potential activation role of genes in Set 2 on genes in Set 1 could be further tested in future experiments.

### Unveiling heterogeneity and resistance mechanisms through single-cell RNA sequencing analysis

By analyzing the bulk RNA-seq data of co-cultured cancer and CTLs, we have revealed essential genes that were differentially expressed in OVA cancer cells and antigen-specific CTLs, respectively. On the other hand, we also observed from the time-lapse images that some cancer cells survived longer and some CTLs stayed longer on cancer cells. To further investigate the molecular origins of this heterogeneity within each cell type, we performed single-cell RNA sequencing (scRNA-seq) on the sorted cancer cells and CTLs.

First, we applied the scRNA-seq clustering of all cells at a resolution of 0.5, and then projected the results on the UMAP (see **Methods**). Since we sorted the cancer cells and CTLs before scRNA-seq, we could display four groups of cells under the two experimental conditions on the UMAP: WT-cancer, CTLs co-cultured with WT-cancer (WT-CTLs), OVA-cancer, and CTLs co-cultured with OVA-cancer (OVA-CTLs) (**Figure 5A**). We found nine sub-clusters of cells in total (**Figure 5A**). Interestingly, there was a distinct cluster of CTLs (C9) associated with sorted OVA cancer cells but not with WT ones (**Figure 5A**). With 9 clusters of cells identified, we conducted differential expression gene (DEGs) analysis and generated a heatmap to provide an overview of the differences among the various clusters (see **Methods** and **Fig. S15A**). For cancer cells, consistent with the bulk RNA-seq data, the gene expression patterns of the WT cancer-cell clusters (C1 and C2) were significantly different from those of the OVA cancer-cells (C3 and C4) (**Fig. S15A**). For CTLs, there were no significant differences in the gene expression pattern among the other T-cell clusters (C5 to C8) compared with C9 (**Fig. S15A**).

**Figure 5.**
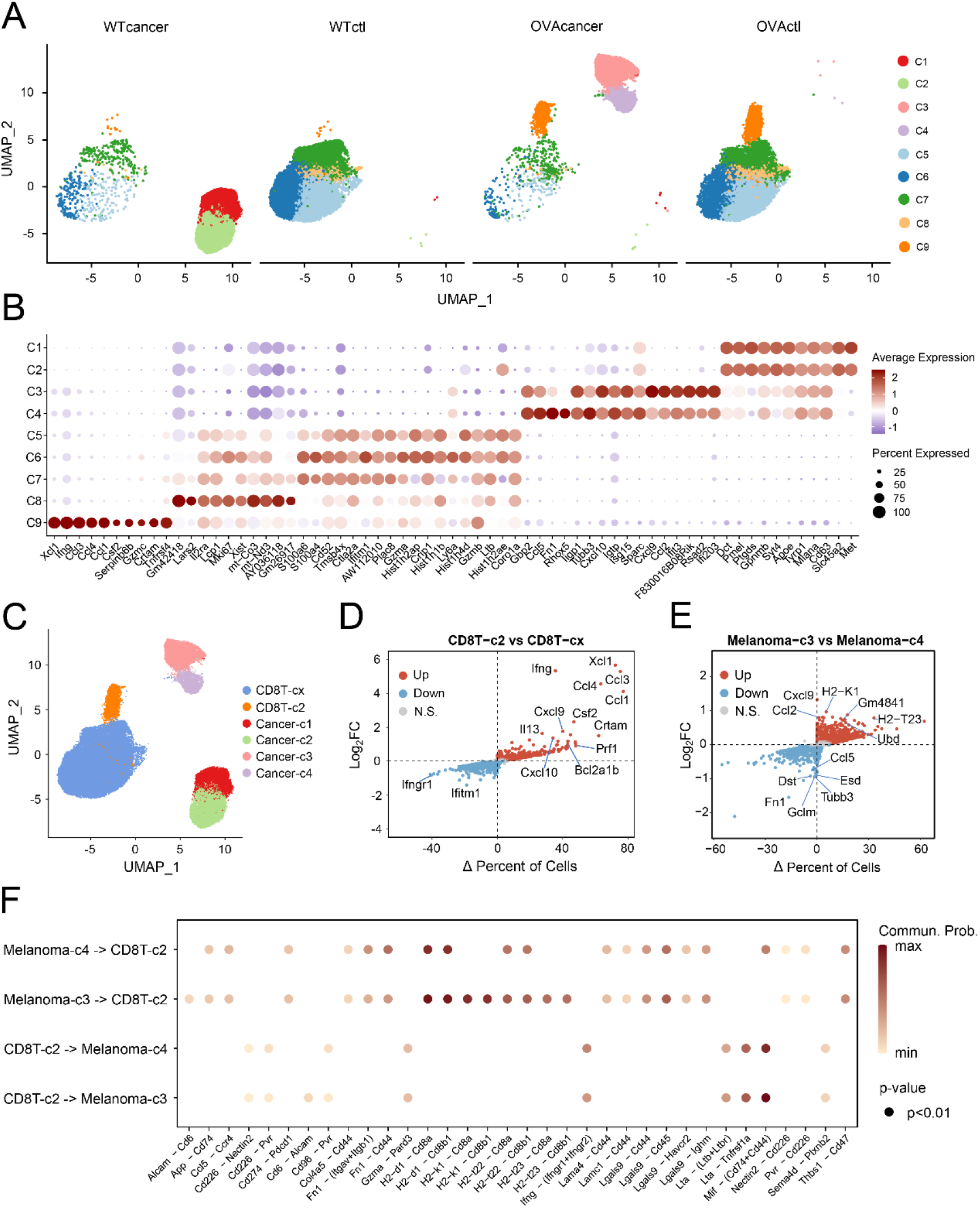
Analysis of the single-cell RNA sequencing data of cancer cells and CTLs. *(A)* UMAP of scRNA-seq data of the sorted cancer and T cells. All cells were combined to perform the clustering, and 9 cell clusters (C1 to C9) were identified after removing ones with low RNA contents or high mitochondrial RNAs (see **Methods**). WTcancer and WTctl (OVAcancer and OVActl) cells were flow cytometry-sorted following 12 hours of co-culture, with EpCam and CD8 as the marker, respectively (see **Methods**). *(B)* Top ten differential expressed genes of the cell clusters shown in panel *A*. The color represents the average expression of cells in the indicated cell cluster. The size of the circle represents the percentage of cells that express the DEG in the indicated cell cluster (see **Methods**). *(C)* Merge of non-unique CTL-subclusters and rename of cancer and CTL clusters. OVActl cells belonging to CTL-subclusters that overlapped with the ones in WTctls were merged into one new subcluster named as CD8T-cx. *(D)* Gene-expression differences between OVActl cells that belong to CD8T-c2 and CD8T-cx in panel *C*. Genes expressed in more cells (x-axis) from CD8T-c2 and CD8T-cx as well as a higher average expression (y-axis) were shown in red and blue, respectively. *(E)* Gene-expression differences between Cancer-c3 and Cancer-c4 cells in panel *C*. Genes expressed in more cells (x-axis) from Cancer-c3 and Cancer-c4 as well as a higher average expression (y-axis) were shown in red and blue, respectively. *(F)* Ligand-receptor cellular interactions between CTLs (CD8T-c2) and cancer cells (Cancer-c4 or -c3) in the coculture of OVA cancer cells and CTLs (see **Methods**). Only the interactions with a *p* value smaller than 0.01 were displayed. The color represents the communication probability as indicated by the color bar.

Second, we investigated the differentially expressed genes between sub-clusters shown in **Figure 5A** (see **Methods**). For WT-CTLs, the top 10 signature genes of the subcluster C9 (**Figure 5B**) were all in the up-regulated gene list from the bulk RNA-seq data (**Figure 4A**). Interestingly, these genes had significantly lower expression in the other OVA-CTL sub-clusters. This result suggested that only a subset of OVA-CTLs contributed significantly to the difference with WT-CTLs in the bulk RNA-seq results. In addition, by performing a volcano analysis (see **Methods**) between CTLs in C9 (CD8T-c2 in **Figure 5C**) and the merged cluster of C5 to C8 (CD8T-cx in **Figure 5C**), we found that the CTLs in CD8T-c2 not only had higher expression of chemokines (e.g., Xcl1, Ccl1, Ccl3, and Ccl4), adhesion proteins (e.g., Crtam) and effector proteins (e.g., Ifng and Prf1), but also had a higher percentage of cells expressing these proteins (**Figure 5D** and **Dataset S5**).

For cancer cells, the scRNA-seq data analysis also revealed the heterogeneity within OVA-cancer cells in their response to CTLs attack. For example, Fn1, Tubb3 and Dst, which could be responsible for cancer-cell migration and metastasis (27–29), had higher expression in the cells from Cancer-c4 than those from Cancer-c3 (**Dataset S5**). In the volcano plot between Cancer-c4 and Cancer-c3 (**Figure 5E**), we showed that Fn1 had a higher expression in Cancer-c4 and 15% more cells in Cancer-c4 than Cancer-c3 expressed Fn1. For Tubb3 and Dst, although the percentage of cells expressing it was the same between Cancer-c4 and Cancer-c3, its average expression level was two-fold higher in Cancer-c4 than that in Cancer-c3 (**Figure 5E**). In addition, the cells in Cancer-c4 had lower expression of T-cell chemokines, such as CXCL9 and CXCL10, than those in Cancer-c3. As shown in **Figure 5E**, the average expression level of CXCL9 was two-fold higher in Cancer-c3 than that in Cancer-c4, though the percentage of cells expressing CXCL9 was very similar. In addition, Cancer-c4 cells expressed less antigen-presentation proteins, such as H2-T23 and H2-K1. These observations suggest that, within the OVA cancer-cell population, some cells (Cancer-c4 in our study) might develop resistance to T cell attack, while others followed the classical response to CTL-attack.

Next, with the differentially expressed genes identified, we applied pathway enrichment analysis to investigate the functional enrichment of these genes. For the CD8T-c2 cluster, the differentially expressed genes (**Dataset S5**) were enriched in pathways related to positive regulation of cytokine production, cell killing, regulation of cell-cell adhesion, *etc.* (**Fig. S15B**). These results were consistent with the bulk RNA-seq analysis. For the Cancer-c3 cluster, the differentially expressed genes (**Dataset S5**) were enriched in pathways related to response to interferon, positive regulation of cytokine production, cytokine-mediated signaling response, *etc.* (**Fig. S15C**). For the Cancer-c4 cluster, the differentially expressed genes (**Dataset S5**) were enriched in pathways related to cell adhesion, ameboid cell migration, regulation of cell morphogenesis, *etc.* (**Fig. S15D and Dataset S5**). A gene-set enrichment analysis focused on epithelial-to-mesenchymal transitions further confirmed that DEGs in the Cancer-c4 cluster are enriched for genes associated with this process (**Fig. S15E**). These results also suggested that OVA cancer cells in the Cancer-c4 cluster might have a higher metastatic potential and stronger resistance towards CTL-attack after interacting with CTLs.

Finally, beyond demonstrating the change in the gene-expression of cancer and T cells after the co-culture, we further investigated the potential ligand-receptor pairs that mediated the interactions among cells. Specifically, we performed CellChat analysis using the scRNA-seq data of cancer and T cells at 12 hours after the co-culture. Notably, Cancer-c3 cells have several unique interactions with the CTLs (**Figure 5F**), such as Cd6-Alcam, H2-k1-Cd8a/Cd8b1 and H2-t23-Cd8a/Cd8b1. Some of these interactions have been shown to be important for the activation/expansion/adhesion of T cells (30, 31) or the antigen-presentation of cancer-cells (32, 33). The lack of these interactions between Cancer-c4 cells and antigen-specific CTLs might indicate that these cancer cells were more resistant to the CTL-attack. Moreover, it was interesting to note that only CTLs in CD8T-cx cluster interacted with OVA cancer cells via CXCL9-CXCR3 and CXCL10-CXCR3, which could be responsible for the motility behavior of antigen-specific T cells toward cancer cells **(Fig. S15F**).

## Discussion

In this work, we used a semi-2D co-culture system to characterize the motility patterns of cytotoxic T cells co-cultured with cancer cells expressing a specific antigen. We found that antigen-specific CTLs had higher directional persistence outside of cancer-cell clusters and longer dwell time on cancer cells than non-specific CTLs. In addition, with the help of computational simulations we showed that higher directional persistence can reduce the first arrival time of CTLs on cancer cells, while longer dwell time can increase the accumulation of CTLs on cancer-cell clusters. These motility patterns of CTLs might help them to locate and collectively unleash the attack on cancer cells.

To decipher the potential mechanisms underlying the observed motility patterns of CTLs, we also performed gene expression analysis and found that both cancer cells and CTLs expressed more chemokines and cytokines after the attack of CTLs. These molecules might increase the directional persistence and bias of CTLs towards cancer-cell clusters. Further knock-out experiments will be required to quantify the effects of these molecules on CTL motility. Previous studies have shown that CC3/4 should be responsible for attracting CTLs in 3D co-culture systems (34). And they might be responsible for the enhanced directional bias of CTLs in our semi-2D co-culture system.

Furthermore, to understand the longer dwell-time of CTLs on cancer cells, we identified several candidates for the adhesion between cancer cells and antigen-specific CTLs, such as MHC-I/TCR and integrin, based on gene expression data. The effects of these molecules on dwell time should be further consolidated using knock-out experiments in the future.

In addition to CTLs, by studying the scRNA-seq data, we also discovered that cancer cells differentiated into two groups after interacting with antigen-specific CTLs. One group expressed less antigen-presenting molecules and chemokines but more EMT-related and immunosuppressive genes. These observations might help to provide mechanistic insights on the potential immune evasion of cancer cells. It has been suggested that cancer-cell EMT induced by immune attack can suppress the activity of CTLs (35, 36). Detailed mechanistic studies are still required to unveil the detailed biological mechanisms.

Interestingly, we also observed a decreasing pattern of directional persistence as CTLs approached cancer-cell clusters. We hypothesized that this was related to the interactions between CTLs and collagen fibers in the system. Our preliminary analysis shown in supplementary Information suggested that a higher density of collagen fibers might lower the direction persistence. We will do future experimental work to systematically quantify the effects of fiber network on CTL motility.

Finally, we only investigated the interactions between cancer cells and antigen specific CTLs in the semi-2D co-culture. In real tumors, there are other types of cells, such as cancer-associated fibroblasts, endothelial cells, and tumor-associated macrophages. Co-culturing more types of cells *in vitro* hold the promise to help understand the effects of paracrine and autocrine interactions on the spatial distribution of antigen-specific CTLs (37).

## Materials and Methods

### Experimental procedures and manual analysis

#### Cell culture

B16-F10 cells were kindly provided by Stem Cell Bank, Chinese Academy of Sciences. B16-F10 murine melanoma cells were expressed ovalbumin and Ca^2+^ sensor GCaMp6s (B16-OVA) by lentivirus infection. B16 cancer cells were cultured in DMEM (GIBCO) supplemented with 10% FBS (GIBCO) and 1% penicillin/streptomycin (GIBCO).

The CD8^+^ T cells with purity above 95% were obtained from female C57BL/6J-OT-I spleens by positive isolation. The naive OT-1 CD8+ T cells were stimulated by Dynabeads® Mouse T-Activator CD3/CD28 (GIBCO) (38) and cultured in RPMI medium 1640 supplemented with heat inactivated 10% FBS (GIBCO), 1% penicillin/streptomycin (GIBCO), 2mM L-glutamine (GIBCO), 50mM β-mercaptoethanol (GIBCO) and 30 U/ml mouse recombinant IL-2 (Biolegend).

#### Co-culture of cancer and T cells

B16-F10-WT and B16-F10-OVA cells were seeded onto Poly-D-lysine (Sigma) coated µ-Slide 8 well (ibidi) and incubated overnight. B16-F10-OVA cells were subjected to a 1-hour pulse with 1 µM OVA peptide at 37°C, followed by two washes with serum-free DMEM. Subsequently, the collagen I solution (Corning) was adjusted to the desired pH of 7.5 using NaOH and then mixed with 10XDPBS. Activated OT-1 CD8^+^ T cells, labelled with anti-CD45-PE at an E:T (effector/tumor cell) ratio of 2:1, were resuspended in the prepared collagen solution on ice. The resulting mixture was added to the center of the 8 wells. After polymerization for 45 min at 37°C, 400 µL of complete medium was added. The resulting mixture was added to the center of the 8 wells. After polymerization for 45 min at 37°C, 400 µL of complete medium was added.

#### Live-cell imaging-based cytotoxicity assay

Bright field and time-lapse fluorescent images of FITC (480 nm/512-550 nm), TxRed (540 nm/577-632 nm) were captured at a resolution of 2 min/frame for a duration of 24 hours using NIKON TI2-E system with a 20x objective lens. All co-culture experiments were conducted at 37 °C and 5% CO_2_ within an incubation chamber that enclosed both the microscope stage and body.

#### Cell sorting for bulk and single-cell RNA sequencing

After co-culture, a collagenase solution from Roche was added to the µ-Slide 8 well and incubated for 15 minutes to release T cells from the collagen gel, followed by trypsin EDTA for 5 minutes to detach all cancer cells. The cancer cells and T cells in suspension were isolated using Dynabeads positive selection with EpCam antibody for cancer cells and CD8 antibody for T cells. The bead-bound cells were then resuspended in a release buffer and the bead-free cell suspension was transferred to new tubes for further bulk and single-cell RNA sequencing.

#### Manual analysis of CTL recruitment, division time and interaction time

For measurement of effector cell recruitment (**Figure 1C**), the detected regions of cancer cells were expanded by adding a belt area with a constant diameter of 15 μm (“dilation circle”), and CTLs within the dilated region were manually counted. Division time and interaction time of OT-1 CTLs were analyzed by ImageJ (**Figure 1E&1G**). Migration velocity was processed using Imaris software (RRID:SCR_007370) (**Figure 1F**).

#### Quantification and computational modelling Quantification of CTL trajectories

CTLs were identified using the TxRed fluorescence channel and their trajectories were generated utilizing Trackmate 7 (39), an open-source Fiji plugin. First, the T cell spots were detected using the LOG (Laplacian of Gaussian) detector. The estimated object diameter was set to 14 μm and the quality threshold was set to 0.18. Subsequently, track segments were established using the LAP tracker, with the maximal distance allowed for frame-to-frame linking set to 25 μm. Additionally, only tracks with a duration of at least 10 frames were selected for further analysis. Finally, the resulting track table and spot table were obtained. The former provides information about the trajectories of the CTLs over time while the latter contains the coordinates of all T-cell centroids.

#### Identification of cancer-cell clusters

Cancer-cell clusters were imaged by splitting the FITC fluorescence channel. To extract the main tumor region, basic image processing was conducted using Fiji (ImageJ v.1.54f). This encompassed threshold adjustment, Gaussian blur, and background subtraction. Subsequently, the preprocessed tumor images underwent further processed using customized algorithms in the MATLAB Image Processing Toolbox. These operations included morphological closing, hole filling, edge smoothing, and the removal of small impurities. Cancer-cell clusters were defined as connected regions with an area exceeding 400 pixels. These operations were then applied to stacks of tumor images, generating a time sequence of binarized cancer-cell clusters data.

#### Quantification of the density of CTLs

To quantify the accumulation of CTLs on cancer-cell clusters over time (**Figure 2C**), the density of CTLs on cancer-cell clusters was calculated by dividing the number of CTLs present on cancer-cell clusters by the area of tumor regions, measured in cells/mm^2^. The positions of CTLs obtained from Trackmate tracking relative to cancer-cell clusters were determined by assessing whether the centroids of CTLs overlapped with tumor regions. The curves were smoothed using a coarse rate of 0.05.

The density of CTLs around the cancer clusters (cells/mm^2^, **Fig. S2**) was calculated by dividing the number of T cells within each dilation belt (enlarged to a diameter of 5 μm) by the area of the dilation belt, which expanded every 5 μm from the boundary of the cancer clusters, covering a total range of 0∼45 μm. Additionally, the density of T cells on cancer clusters was also included. A 3D representation of CTL density around cancer clusters was displayed for 6 h, 9 h and 12 h, respectively.

The total density of CTLs within the imaging field (**Fig. S13**) was quantified by dividing the total number of CTLs in the imaging field by the total area of the field, also measured in cells/mm^2^. The curves were smoothed using a coarse rate of 0.05.

#### Dwell-time distribution of CTLs on cancer-cell clusters

To quantify the duration of CTL interactions with OVA cancer cell and WT cancer cell, the dwell-time distribution of CTLs was assessed (**Figure 2D, Fig. S4 & Fig. S5**). Firstly, a dwelling event was defined as the occurrence when the centroid of CTL was within a minimal distance of less than 5 μm from the boundary of cancer-cell clusters. This criterion was established based on radius of the CTL is approximately 5∼6 μm. The dwell-time was then calculated by determining the number of consecutive frames in which CTL was present on cancer-cell clusters. In cases where a CTL attached to the tumor cell clusters multiple times within a track, each attachment was considered as a separate dependent adhesion event.

To compare the dwell time distributions between CTLs cocultured with OVA cancer cell and those with WT cancer cell within different stages, the adhesive time distributions during three specific time periods were statistically calculated: 0-6 hours, 6-12 hours, and 12-18 hours.

#### Directional bias and persistence analysis

To explore the motility patterns of CTLs with respect to their distance towards cancer cell clusters in the two co-culture systems, we conducted a study utilizing the concepts of directional bias and persistence as introduced by Weavers *et al.*(25). Directional bias, denoted by 𝛼, represents the angle between the instantaneous velocity of CTLs and the direction from CTLs to the nearest point on the cancer-cell cluster, reflecting the ability of cells to move in the direction of an attractant source. Additionally, directional persistence, denoted by 𝛽, measures the angle between CTL velocities in two consecutive steps, indicating the ability of cells to maintain their movement direction. The bias angle distributions range from 0° to 180° whereas the persistence angle distributions range from −180° to 180°. The cosine function was applied to the angles to normalize the values between −1 and 1, where a higher value indicates a greater bias or persistence.

For each CTL spot, the distance to the nearest pixel of the cancer cluster boundary was calculated. The trajectory segments of CTLs located outside the cancer clusters were exclusively selected for analyzing directional bias and persistence. These segments were then grouped based on their distances from the nearest cancer-cluster boundary, with each group defined by 5 μm intervals. Subsequently, the averaged bias or persistence was computed for each distance-interval. To compare the behavioral dynamics of CTLs, the average bias and persistence distributions for CTLs cocultured with OVA cancer cells and those with WT cancer cells were calculated over three specific time-periods: 0∼6 h, 6∼12 h, and 12∼18 h, respectively (**Figure 2F&2G**). Additionally, analyses were conducted per 3-hour intervals, as shown in the supplementary information (**Figure 2G&2H**).

#### Mean square displacement of CTLs

To investigate the motility rules and compared the rules in different time intervals, the mean square displacement was calculated to characterize the diffusive behavior for CTLs. The calculation was performed using the following equation (40):

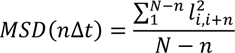

Here, 𝑙_𝑖,𝑖+𝑛_ represents the Euclidean distance of a CTL from frame 𝑖 to frame 𝑖 + 𝑛; *N* represents overall time periods for calculation; *n* represents the time lag.

The MSD can also be described using a power-law form (41):

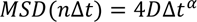

The trajectory segments of CTLs before reaching the cancer cell clusters were extracted. Then, the logarithm of the MSD as a function of the time was fitted, and the linear portions of the curve were identified (**Fig. S7**). To ensure credibility and accuracy of the calculations, only the first 10 frames of continuous segments in the tracks were selected for the fitting process.

#### Quantification of collagen fibers

To investigate potential variations in collagen fiber distribution and arrangement between WT and OVA cancer cell environments, collagen fiber density and orientation were analyzed around cancer clusters in two co-culture systems at 0 h and 6 h, respectively.

For the analysis of fiber orientation, CT-fire software (42) was utilized to extract and calculate the directional alignment angles between collagen fibers and cancer boundaries. (**Fig. S10**).

For the assessment of fiber density, a threshold was manually set to generate binarized images of collagen fibers. Subsequently, fiber density was determined at varying distances from cancer clusters, by dividing the number of fiber pixels by the area of circular rings extending at uniform 5 μm intervals. To characterize the distribution around the cancer clusters and reduce the error, the average fiber density from Z1 to Z5 was calculated (**Fig. S11**).

#### Correlation test among directional bias, directional persistence and step size

To investigate whether there exists correlation among directional bias, directional persistence and step size, Spearman correlation tests were performed by per 5 μm within 0∼6 h, 6∼12 h, and 12∼18 h, respectively (**Fig. S12**).

#### Agent-based models

To further investigate the impact of observed differences in T-cell motility patterns on their searching efficiency and the accumulation on cancer-cell clusters, we constructed 2D agent-based models that incorporated four key features of T cells: dwell time, directional bias, directional persistence and step size. In addition, computational simulations offer the flexibility to manipulate the initial distributions of CTLs and cancer clusters, which is challenging to control under experimental conditions. To address this, we developed two models: a single-cancer-cluster model (**see Fig. S13**) and a multi-cancer-cluster model (**Figure 3**), allowing us to explore the potential effects of the initial distribution of CTLs and cancer clusters.

The multiple-cancer-cluster model is based on the following assumptions:

1. The model is implemented on a 2D square region with dimensions of 590 μm*590 μm, matching the experimental scale. The model includes CTLs and cancer-cell clusters. To simplify the modelling process, the models disregard the mobility of cancer cells, as well as the proliferation or death of both CTLs and cancer cells, and the weak correlation among step size, directional bias and persistence.
2. To simplify the model, the initial distribution of cancer-cell clusters is identical to Series 1 in the experiments. Initially, 200 CTLs are randomly dispersed outside cancer-cell clusters within the defined region. The simulation utilized parameters derived from Series 1 in the WT and OVA experimental systems, including step size, directional bias, directional persistence as a function of distances towards cancer-cell clusters, and dwell time distributions within 6∼12 h, as significant differences were observed in the directional bias, persistence, and dwell time distribution of CTLs during this period (**Figure 2D, 2F-2G**). The step size, directional bias and persistence distributions are updated with a resolution of 5 μm distance from the nearest cancer-cell boundary.
3. When T cells are outside of cancer-cell clusters, they undergo a distance-dependent persistent or bias walk.

✓ If we specifically focus on the influence of the differences in the distribution of directional persistence, we can determine the position of T cells at any given time by following the equations below.

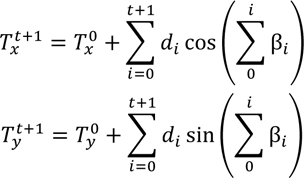

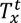 and 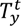 represents the coordinates of a CTL at time t. 𝛽_0_ represents the angle between the initial motion vector and the positive x-axis, which is randomly set. 𝑑_𝑡_ represents the step size from t to t+1, sampled from the step-size distributions. Here, we developed an algorithm to realize the sampling process from a known distribution. 𝛽_𝑡_ represents the turning angles from t to t+1, sampled from persistence distributions based on the distance towards cancer-cell clusters.
✓ Similarly, if we focus solely on the role of directional bias, we can determine the position of CTLs using the following equations. The motion of CTLs depends on searching for the nearest cancer-cluster point at any given time, with the coordinates of denoted as 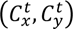.

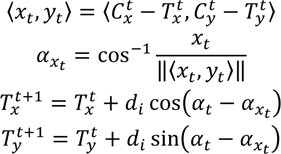

𝛼_𝑥𝑡_ represents the angle between the motion vector and the positive x-axis at time t. 𝛼_𝑡_ represents bias angle from t to t+1 sampled from bias distributions with respect to the distance towards cancer-cell clusters.
4. Once T cells reach cancer-cell clusters, they can remain for a duration time t sampled from the dwell-time distributions. Each CTL underwent 800 computational simulation steps.
5. Then, the effects of the four patterns within two systems were explored individually by altering the directional bias distribution or directional persistence or step-size distribution, while maintaining consistency in the other two patterns (**Table 1**). In terms of searching efficiency, we computed the mean arrival time for all CTLs to initially reach cancer clusters in each simulation, which is depicted as a data point in the violin plot generated using the Violinplot function (https://github.com/bastibe/Violinplot-Matlab) illustrated in **Figure 3D**.
6. In addition, we examined the number of T cells accumulating on cancer-cell clusters over time to explore whether the motion outside of cancer-cell or adhesion could affect the accumulation of CTLs on cancer cell clusters. Each simulation was repeated 100 times. The curves were smoothed using a coarse rate of 0.05.

**Table 1.** Computational modelling to explore the effects of the four pattens.

| Simulation | Step size | Persistence | Bias | Dwell time |
| --- | --- | --- | --- | --- |
| 1 | WT | - | - | - |
|  | OVA | - | - | - |
| 2 | - | WT | - | - |
|  | - | OVA | - | - |
| 3 | - | - | WT | - |
|  | - | - | OVA | - |
| 4 | - | - | - | WT |
|  | - | - | - | OVA |

#### Statistics and reproducibility

Statistical analysis was conducted using MATLAB, and the results are represented as mean±s.e.m. unless stated otherwise. Two-tailed unpaired *t*-tests were performed between two systems. For all figures concerned, the annotations for *p*-value mean that: ns: p>=0.05; *: 0.01<*p*<0.05; **: 0.001<*p*≤0.01; ***: *p*≤0.001. To ensure reproducibility, we conducted five parallel experiments and performed statistics analysis on these experimental studies (**Figure 2**). The computational simulation was repeated 100 times (**Figure 3**).

### RNA sequencing pipelines

#### Bulk RNA-seq data processing

The raw sequencing reads were trimmed to remove sequences with low quality and short read lengths using Fastp (version 0.20.0). Subsequently, HISAT2 was employed to map the high-quality reads onto the mouse reference genome mm10. The read count matrix corresponding to the genes and samples was generated using featureCounts. Next, we transformed the count matrix into an RPKM matrix to qualify the expression levels with in-house scripts. Additionally, the count matrix was subjected to the R package DESeq2 to identify differentially expressed genes with a log2(fold change) cutoff of 1 and a false discovery rate (FDR) of 0.05 (**Figure 4A&4E**). GO term enrichment analysis (MF, molecular function, CC, cellular component, BP, biological process) and KEGG pathway analysis were performed using the R package clusterProfiler (43) based on the up-regulated and down-regulated genes, with significance determined by an FDR threshold of less than 0.05 (**Figure 4C, 4D, 4G&4H**). Gene set enrichment analysis (GSEA) was conducted for GO, KEGG and HALLMARK gene sets using the R package GSEA (**Fig. S14B**). To explore regulatory pairs using the temporal dynamics of gene expression, we employed the AnimalTFDB4 database to extract transcription factors (TFs) in a gene set, and then searched for potential targets of the selected TFs in the TRRUST2 database.

#### ScRNA-seq data processing

Raw gene expression matrices were generated for each sample using the Cell Ranger (v6.1.2) Pipeline coupled with the mouse reference genome version mm10-2020-A. The output filtered gene expression matrices were analyzed using R software (version 4.1.2) with the Seurat package (version 4.1.1) (44). A custom R script was used to combine the expression data and metadata from all libraries corresponding to a single batch. The expression data matrix was loaded into a Seurat object along with the library metadata for downstream processing. The percentage of mitochondrial transcripts for each cell (percent.mt) was calculated and added as metadata to the Seurat object. Cells were further filtered before dimensionality reduction (nFeature_RNA>200 & percent.mt<10%). Expression values were then scaled to 10,000 transcripts per cell and log-transformed (NormalizeData function, normalization.method = “LogNormalize”, scale.factor = 10000). We calculated variable features using the FindVariableFeatures function with selection.method = “vst” and nfeatures = 2000. Effects of variables (percent.mt) was estimated and regressed out using the ScaleData function (vars.to.regress = “percent.mt”), and the scaled and centred residuals were used for dimensionality reduction and clustering.

#### Dimensionality Reduction

To reduce the dimensionality of the dataset, the RunPCA function was conducted with default parameters on linear-transformation scaled data generated by the ScaleData function. Next, the ElbowPlot and DimHeatmap functions were used to identify proper dimensions of the dataset (first 11 PCs).

#### Cell Clustering and Cluster Identification

We first performed the FindNeighbors function before clustering, which takes as input the previously defined dimensionality of the dataset (first 11 PCs). Clustering was performed at varying resolution values using the FindClusters function, and we chose a value of 0.7 for the resolution parameter for the initial stage of clustering. After non-linear dimensional reduction and projection of all cells into two-dimensional space by UMAP, clusters were assigned preliminary identities based on the expression of combinations of known marker genes for major cell classes and types.

#### Sub-Clustering and Secondary Clustering

We filtered out low quality clusters and mixing cell clusters (C1, C6, C9, C16, C17). Sub-Clustering was performed on the two major cell types (CD8 T Cells and Melanoma) data subset to resolve additional cell types and subtypes. Secondary Clustering was performed using a standard workflow similar to before but with different dimensionality (first 12 PCs) and a value of 0.5 for the resolution parameter. We filtered out three clusters (C9, C10, C11), renamed all clusters from C1 to C9 (as shown in **Figure 5A**) and then performed other downstream analyses.

#### Differential Expression Genes (DEGs) Identification and Functional Enrichment

Differential gene expression testing was performed using the FindAllMarkers function in Seurat with the parameter only.pos = TRUE. DEGs were filtered using a minimum log2(fold change) of 0.5, a minimum expression percentage value of 0.3 and a maximal *p* value of 0.05. Dot plots showing the relative average expression of top 10 DEGs in subclusters were visualized (**Figure 5B**). All DEGs’ relative average expression in subclusters was visualized using the DoHeatmap function in Seurat (**Fig. S15A**). GO (BP) enrichment analysis for the functions of the DEGs was conducted using the clusterProfiler (43) (version 4.2.2) R package (**Fig. S15B-D**).

#### DEGs analysis between two groups

A Wilcoxon rank-sum test for differential gene expression comparing CD8T-c2 (C9) versus CD8T-cx (merged clusters of C5 to C8, **Figure 5C**) in OVActl was performed using the FindAllMarkers function (min.cells.group = 0, min.pct = 0, logfc.threshold = -Inf) in Seurat (**Figure 5D**). Differential gene expression comparing Melanoma_c3_Cxcl9 versus Melanoma_c4_Rhox5 (cell type of secondary-clustered dataset) was also performed with the same function and parameters (**Figure 5E**). Percentage difference (Δ Percent of Cells) and log2(fold change) are shown in a volcano plot. Genes highlighted in red or blue have adjusted *p* values < 0.05.

#### Cell-cell communication analysis (CellChat)

We used CellChat (45) (v1.4.0) to infer cell-cell communication based on prior ligand-receptor interaction databases. We separately loaded the normalized counts data of WT group (WTcancer, WTctl) and OVA group (OVAcancer, OVActl) extracted from Seurat object into CellChat and followed the workflow recommended in CellChat to infer and visualize cell-cell communication network. The database of known receptor-ligand pairs was applied to assess cell-cell communication in the selected target clusters (CD8T-c2, CD8T-cx, Melanoma_c3_Cxcl9, Melanoma_c4_Rhox5) of OVA group (**Figure 5F & Fig. S15F**). Interactions were trimmed based on significant sites with *p*<0.01.

## Data and Software Availability

The data and customized codes are available on figshare (https://figshare.com/articles/online_resource/Quantitative_effects_of_co-culture_on_T_cell_motility_and_cancer-T_cell_interactions/25367845). All data needed to evaluate the conclusions in the paper are present in the paper and/or SI Appendix.

## Acknowledgments

This work was supported by the National Key R&D Program of China (2021YFA0911100), the National Natural Science Foundation of China (32170672, 12274308, 32000886, 12004271), Key-Area Research and Development Program of Guangdong Province (21202107221900001), the Guangdong Basic and Applied Basic Research Foundation (2021A1515012461, 2019A1515110186), and Shenzhen Science and Technology Program (ZDSYS20220606100606013).

