## Supplementary material for "Quantitative effects of co-culture on T cell motility and cancer-T cell interactions": Dataset

**This PDF file includes:**

Figures S1 to S15
Legends for Movies S1 to S8
Legends for Datasets S1 to S5

**Other supporting materials for this manuscript include the following:**

Movies S1 to S8
Datasets S1 to S5

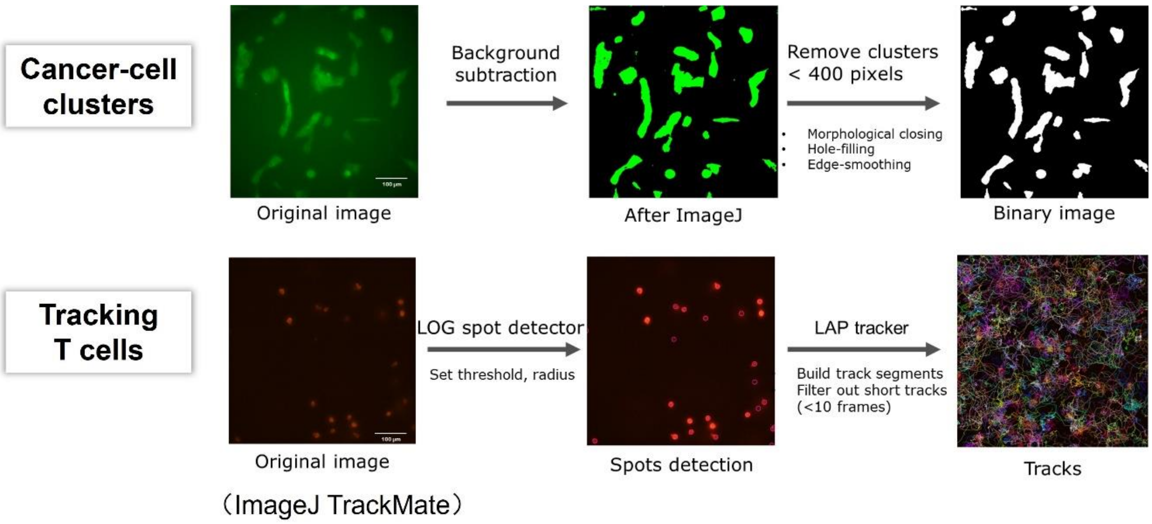


**Fig. S1. Image processing pipelines for cancer-cell clusters and tracking T cells.**

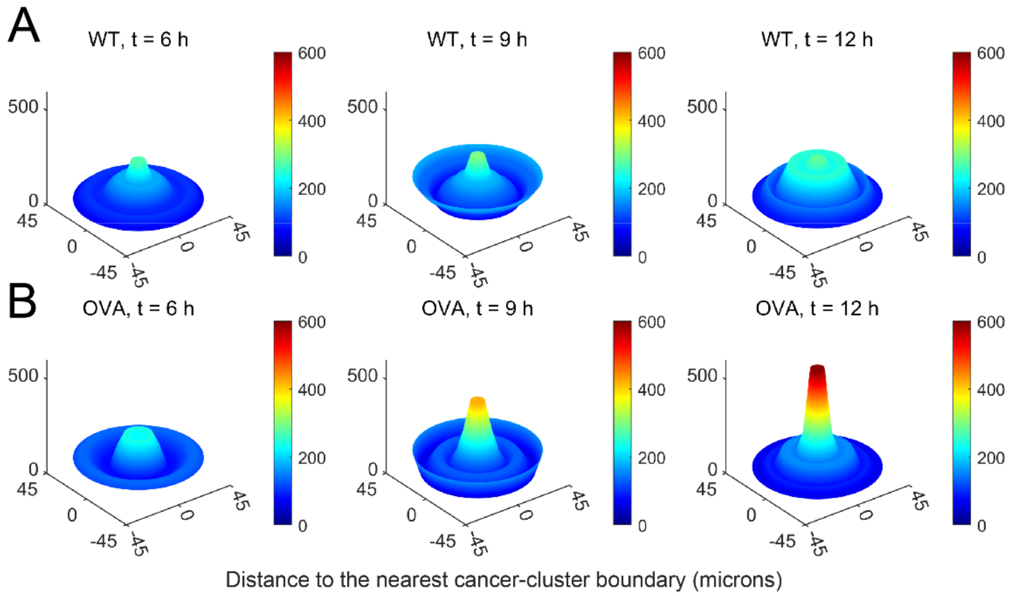


**Fig. S2. Density of CTLs around cancer-cell clusters (cells/mm$^2$) after co-culturing for 6, 9, 12 hours, respectively.** CTLs co-cultured with OVA cancer cells exhibited a significant accumulation around cancer clusters compared to those co-cultured with WT cancer cells within 6-12 h.

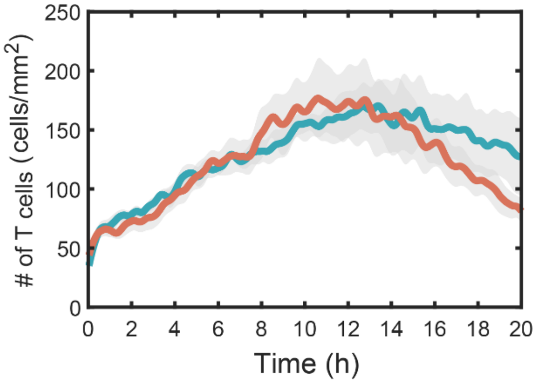


**Fig. S3. Total number of CTLs (cells/mm$^2$) in the imaging field varies over time.** There is no significant difference in the total number of CTLs between the two co-culture systems.

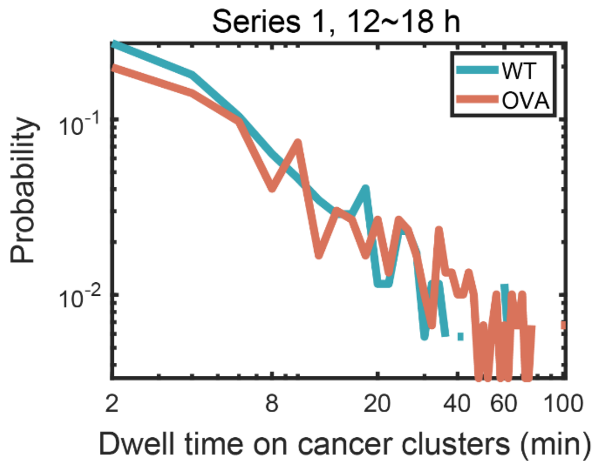


**Fig. S4. Dwell time distribution of CTLs on cancer-cell clusters in Series 1 within 12~18 h.** Similar to Figure 2D (right panel), the distribution on cancer-cell clusters exhibited a longer tail compared to that on the WT cancer-cell clusters.

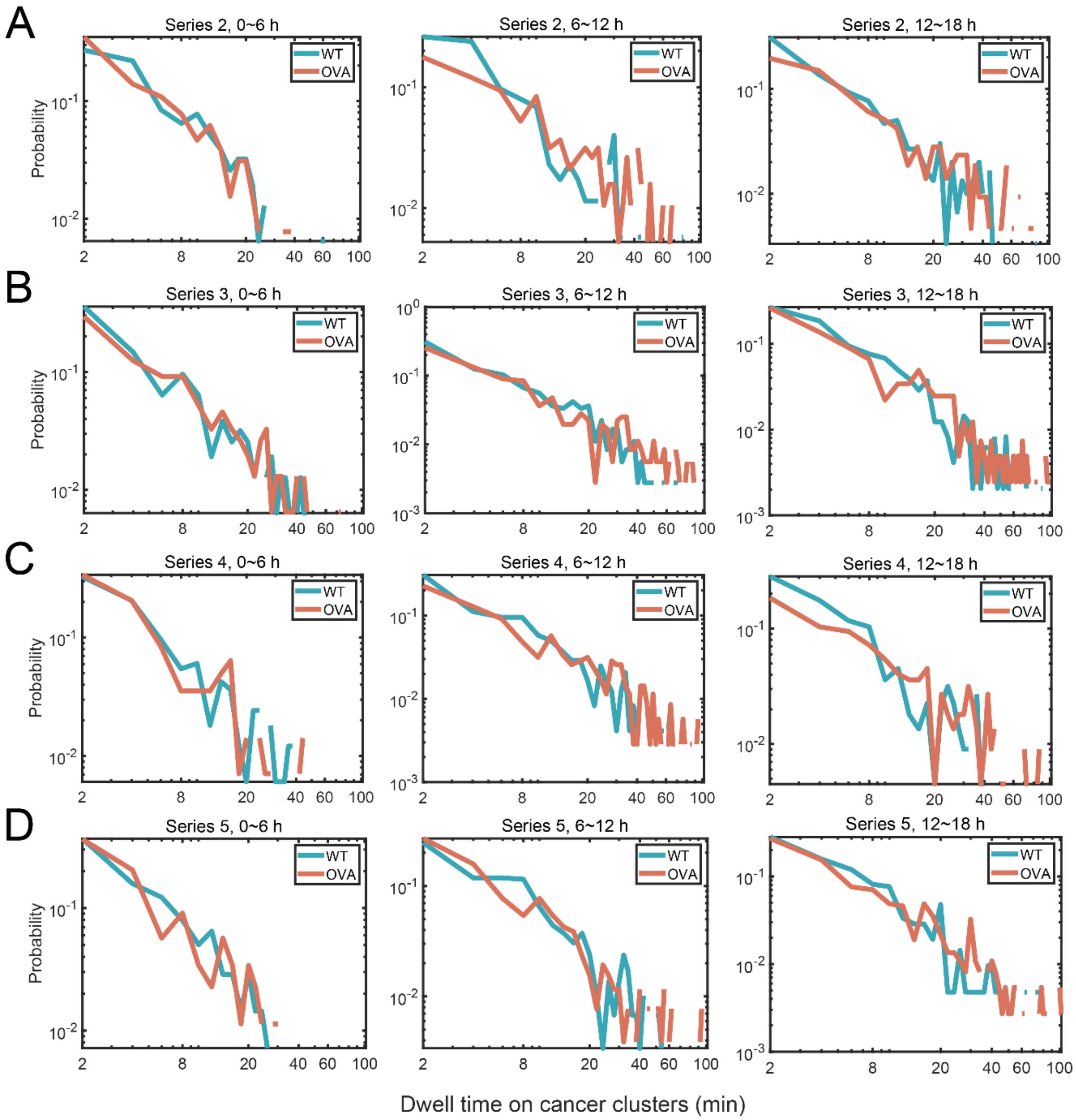


**Fig. S5. Dwell time distribution on cancer clusters across (*A*) Series 2, *(B)* Series 3, *(C)* Series 4 and *(D)* Series 5.** The time intervals from left to right are 0~6 h, 6~12 h and 12~18 h. Similar to the result shown in Series 1 (Figure 2D & Fig. S4), within the first 6 hours, there is no significant difference in the dwell time distribution of CTLs between the two co-culture systems. However, after 6 hours of the co-culture, the distribution on cancer-cell clusters exhibited a longer tail compared to that on the WT cancer-cell clusters. These results are consistent among the five series.

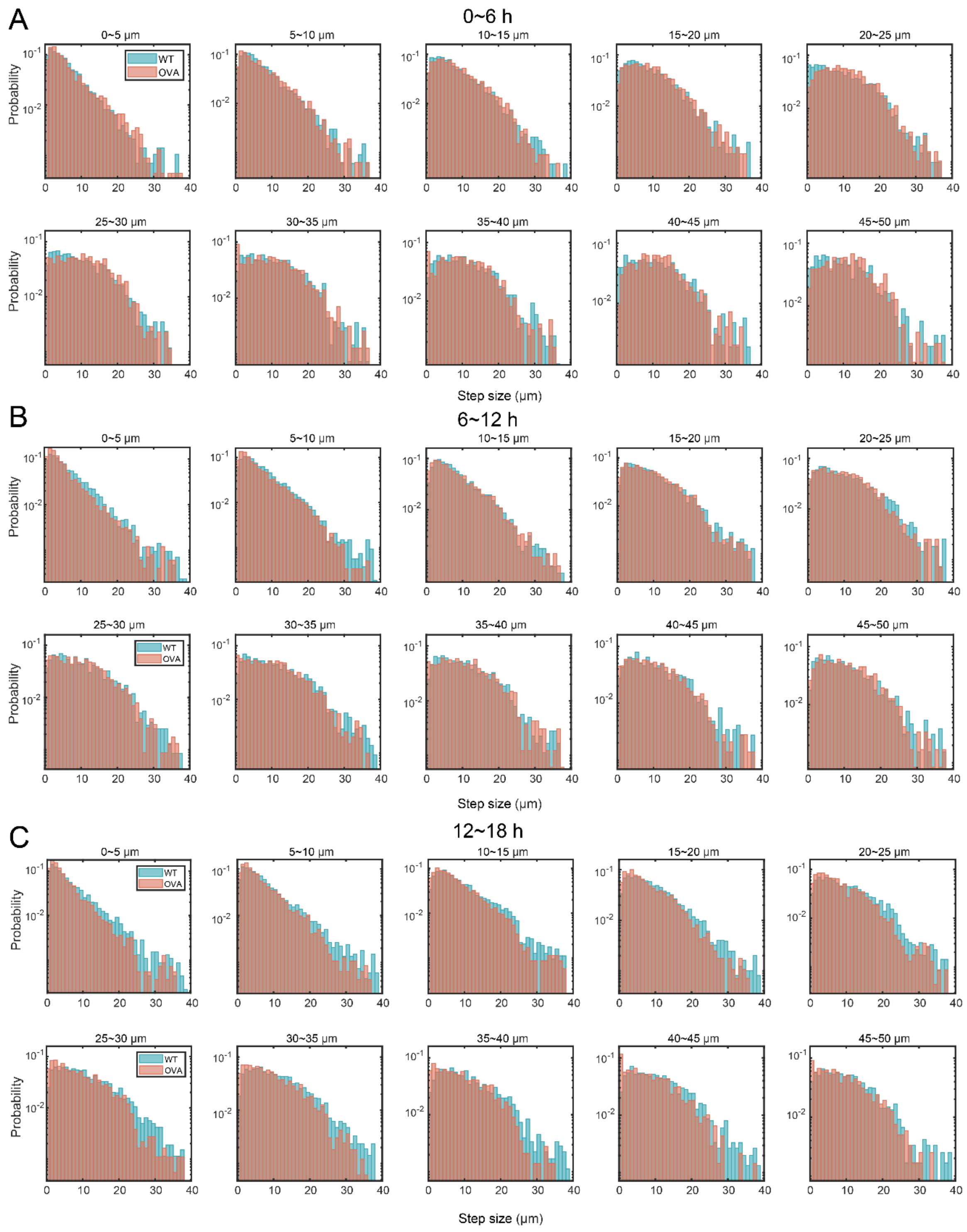


**Fig. S6. Step size distributions of CTLs within 6-hour intervals at each distance from cancer-cell clusters.** The distributions depict CTL step sizes at various distances (per 5 microns) from the cancer-cell cluster within three time-intervals: *(A)* 0~6 h, *(B)* 6~12 h and *(C)* 12~18 h. CTLs within 5 microns from cancer boundary exhibited smaller step sizes, typically less than 3 microns (the first panel of within every time-interval). After 12 h, CTLs co-cultured with OVA cancer-clusters showed shorter step sizes compared to those with WT cancer-clusters.

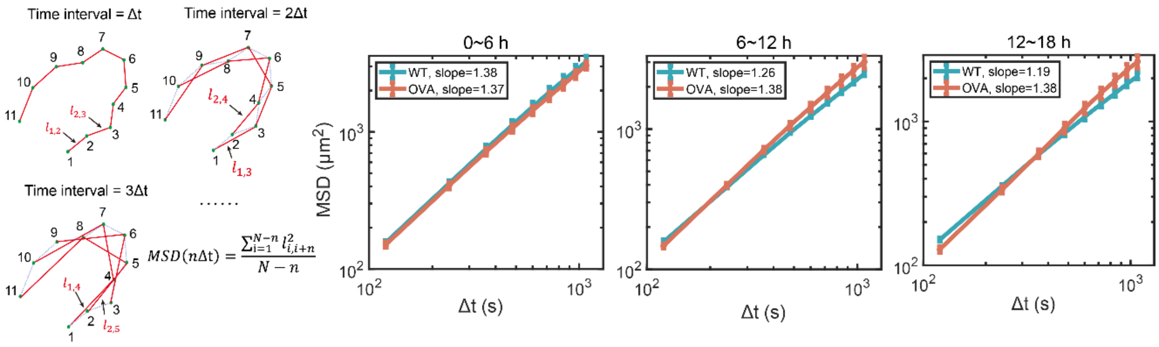


**Fig. S7. Mean square displacement analysis for CTLs within every 6-hour interval. Left, 0-6 h; middle, 6-12 h; right, 12-18 h.** CTLs in both systems demonstrated super-diffusive motion behavior with the exponent α around 1.3. CTLs cocultured with OVA cancer cells exhibited a higher power exponent compared to those cocultured with WT cancer cells after 6 hours.

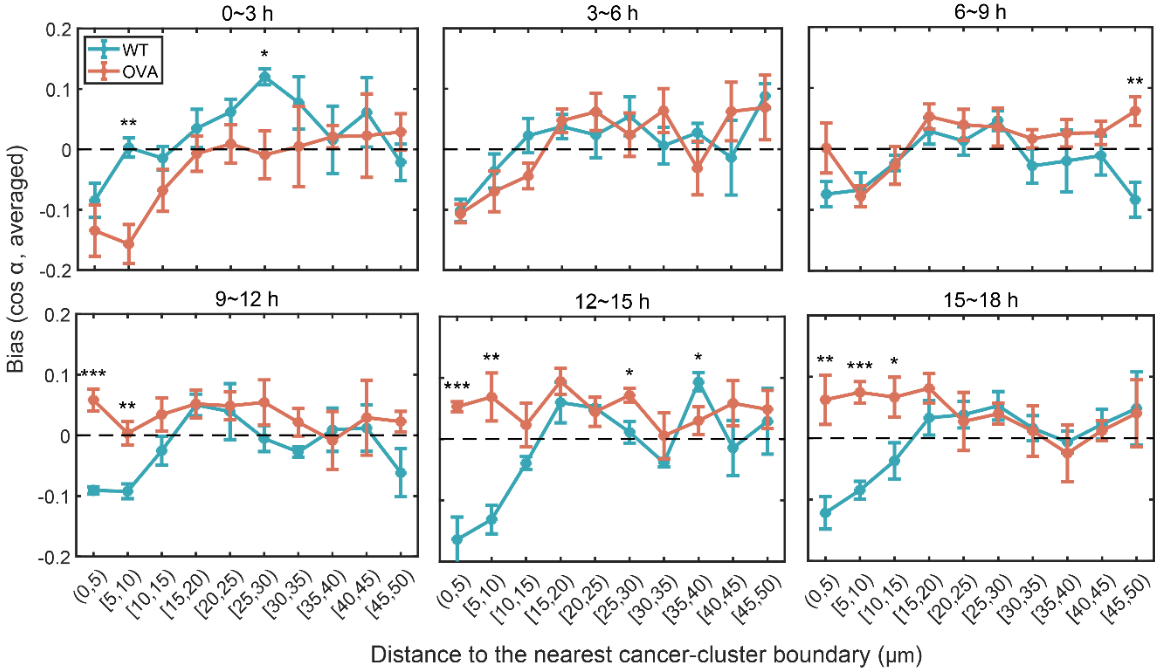


**Fig. S8. Directional bias distributions of CTLs movement towards the nearest cancer cell boundary within each 3-hour interval.** CTLs co-cultured with OVA cancer cells gradually increased their motion bias towards the nearest cancer-cell clusters from the first 6 hours to the 9-18 hours of the co-culture, particularly when they were within 0-10 μm away from cancer-cell clusters.

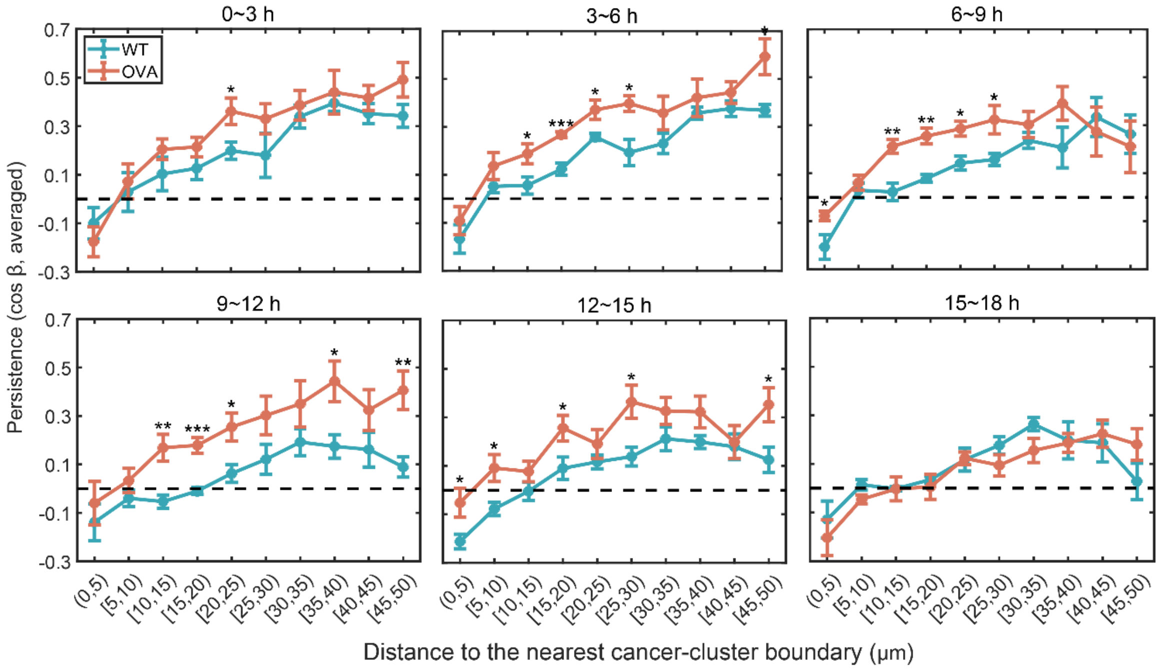


**Fig. S9. Directional persistence distributions of CTLs movement towards the nearest cancer cell boundary within every 3-hour interval.** CTLs in the co-culture with OVA cancer cells displayed considerably stronger persistence compared to those cultured with WT cancer cells, particularly in the range of 10~30 μm away from cancer-clusters during 3-12 h.

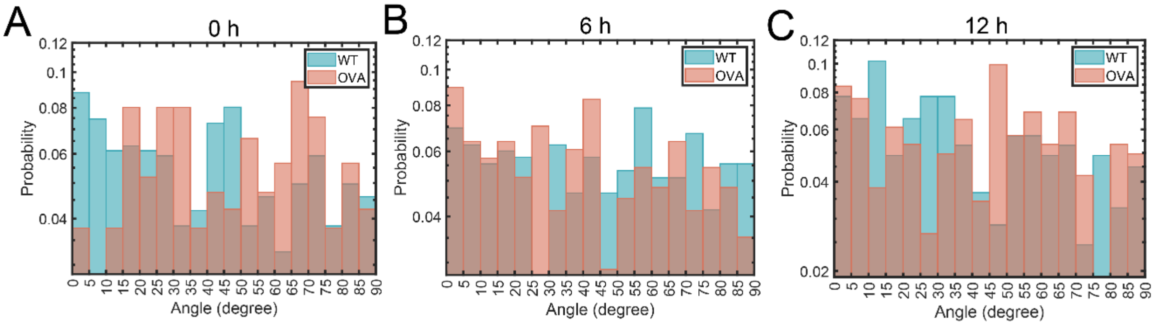


**Fig. S10. The relative angular distribution of collagen fibers to cancer boundaries within 20 microns at 0 h, 6 h and 12 h, respectively.** No significant difference was found between the OVA and WT groups. For each panel, the histogram was generated from a single experiment. The y-axis was transformed to a logarithmic scale.

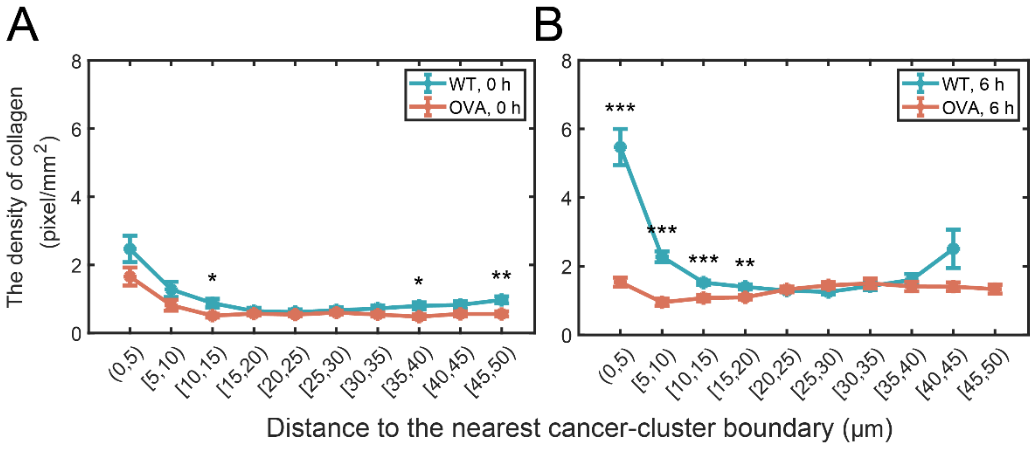


**Fig. S11. Density distribution of collagen fibers around cancer-cell clusters at *(A)* 0 h and *(B)* 6 h.** The results reveal a higher density of collagen fibers in proximity to cancer-cell clusters at both 0 h and 6 h. Compared to OVA group, the density of fibers in WT group were significant higher within a 20 μm distance away from cancer-cell clusters.

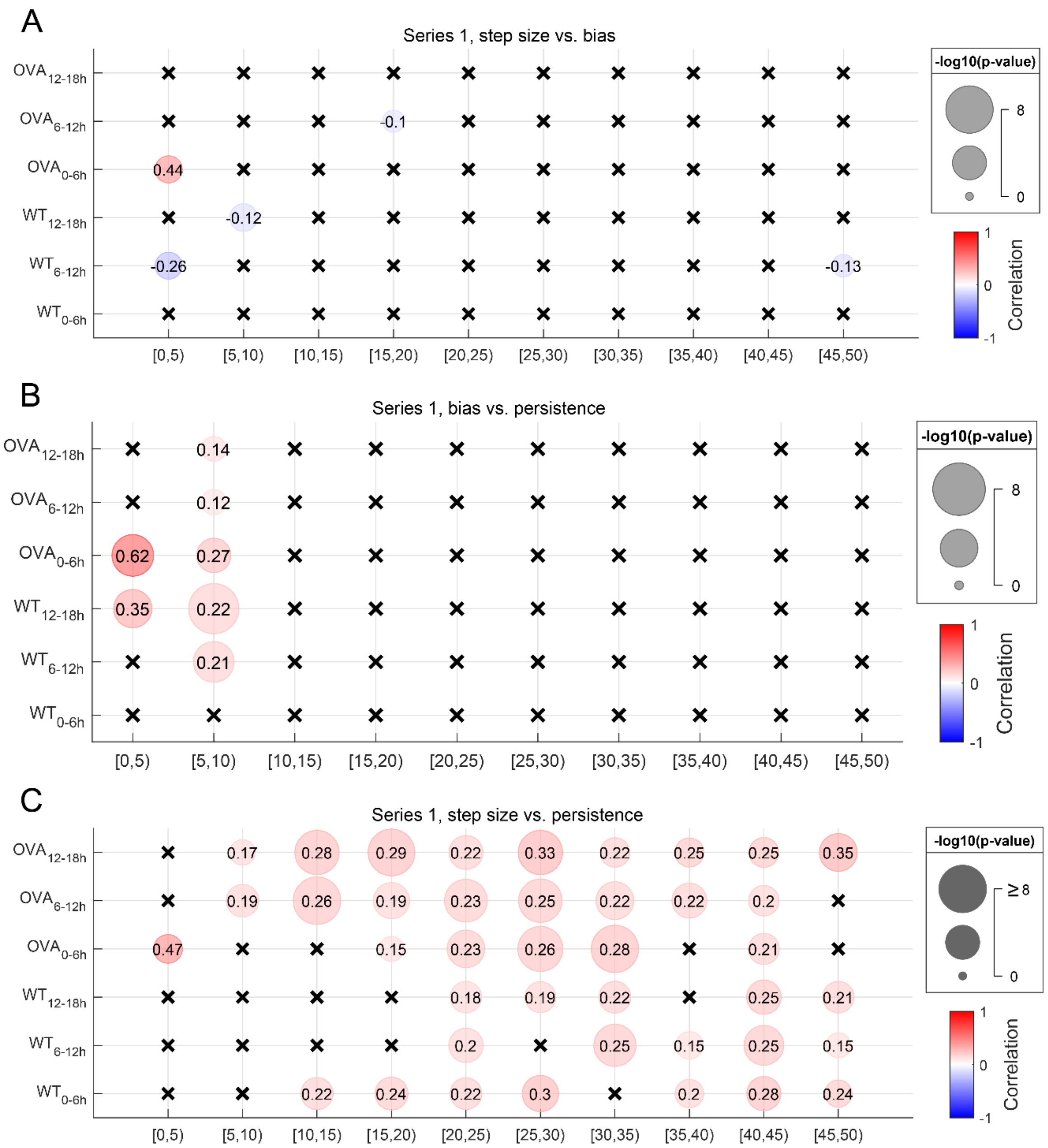


**Fig. S12. Pairwise correlation test among step size, directional persistence and directional bias under different distances.** *(A)* Step size vs. bias. *(B)* Bias vs. persistence. *(C)* Step size vs. persistence. The cross symbol represents $p>0.05$, while the spot symbol represents $p<0.05$. Additionally, the size of the spots corresponds to the *p*-value. Correlation analysis was conducted using the Spearman method.

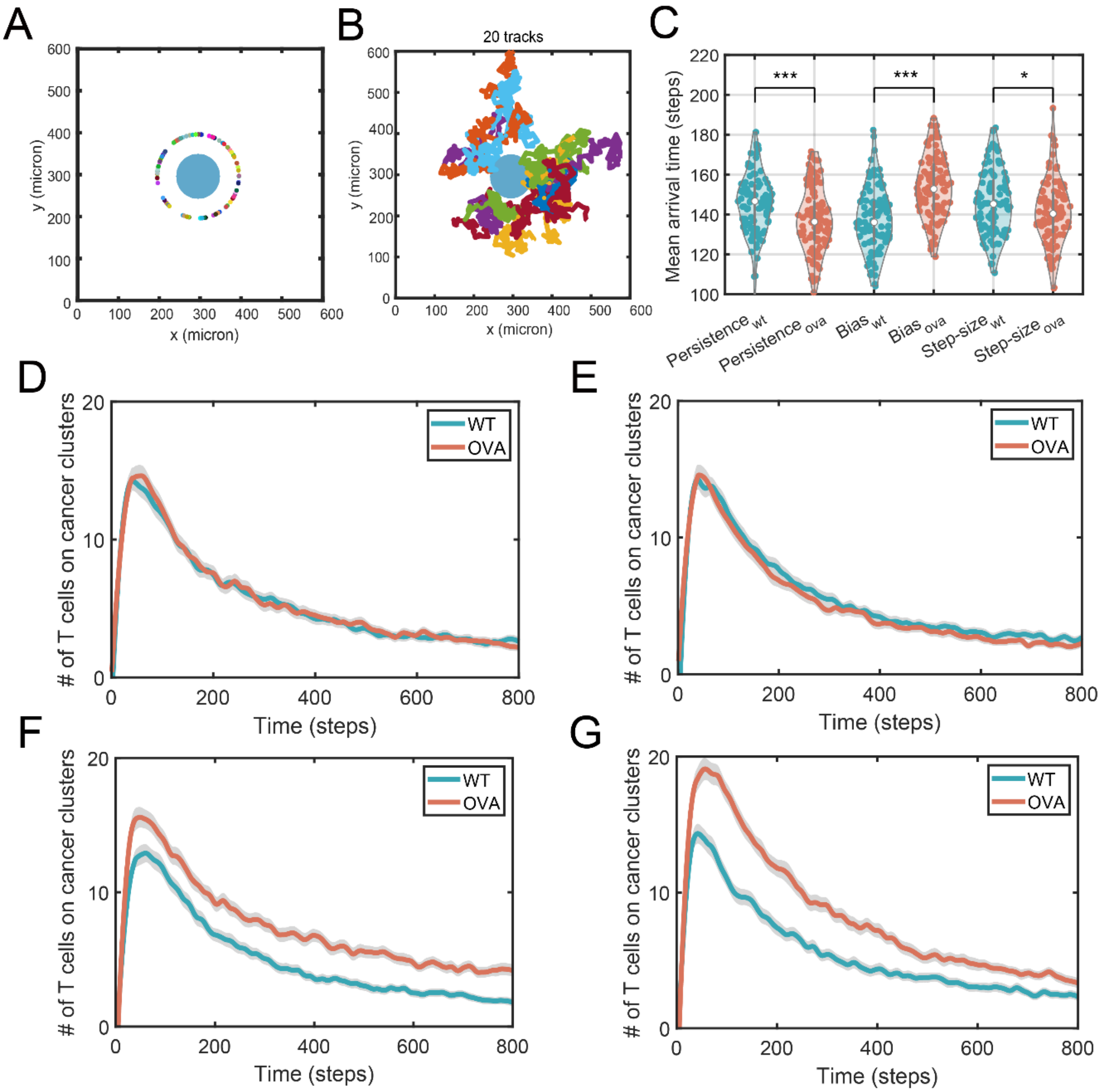


**Fig. S13. Design and data analysis of the single-cancer-cluster model.** *(A)* The schematic representation of single-cancer cluster model. Initially, cancer-cell clusters are distributed within a circle with a 50 μm radius. The center of the cancer is at (295,295), and CTLs are randomly distributed along a circular arc located 100 μm from the center of the cancer cluster. *(B)* An illustration depicting the trajectories of 20 CTLs after moving 100 steps. Each spot represents a CTL. *(C)* Violin plot and statistical significance test of the mean arrival time of CTLs to cancer clusters under different patterns when considering only the differences in the directional persistence or directional bias or step sizes. 100 replicated simulations were performed, with each simulation consisting of 200 cells moving for 800 steps. '*' represents $p$<0.05, '**'represents $p$<0.01 and '***' represents $p$<0.001. *(D-G)*. The accumulation of CTLs on cancer-cell clusters as a function of time when considering different patterns: (*D*) step size, (*E*) directional persistence, (*F*) directional bias, and (*G*) dwell time over time. The shaded area represents the 95% confidence interval of the simulated data, while the bold line represents the mean.

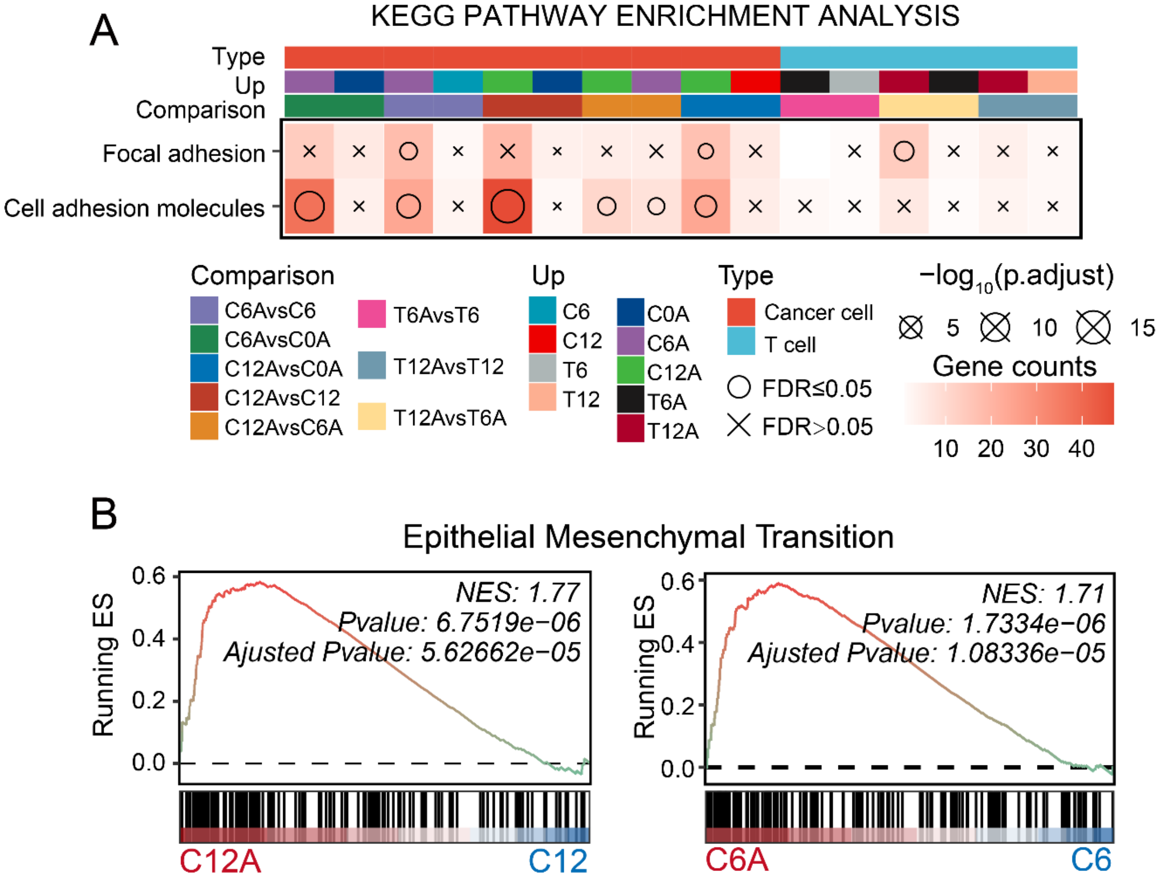


**Fig. S14. Differential gene expression enrichment analysis for bulk RNA sequencing.** *(A)* KEGG pathway analysis of OVA cancer cells, WT cancer cells and co-cultured T cells at 0 h, 6 h and 12 h, respectively. *(B)* Gene Set Enrichment Analysis (GSEA) for hallmark gene sets of EMT between OVA and WT cancer cells at 12 h (C12A vs. C12) and 6 h (C6A vs. C6), respectively.

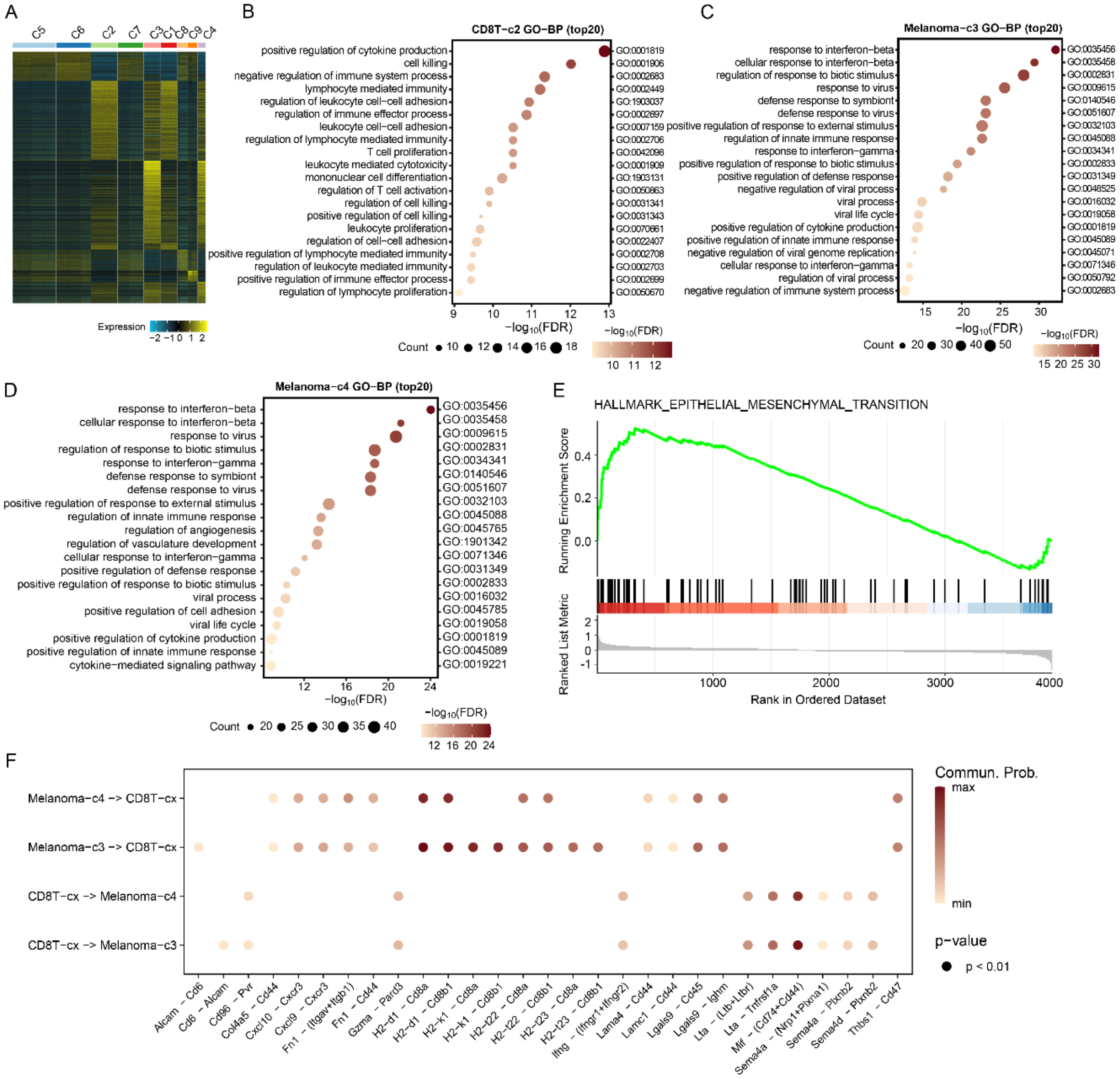


**Fig. S15. Differential gene expression analysis for single-cell RNA sequencing.** *(A)* Heatmap of differential expression genes of clusters from C1 to C9. *(B)* Results of the GO-BP enrichment analysis for the top 20 differentially expressed genes within CD8T-c2 cluster. *(C)* Results of the GO-BP enrichment analysis for the top 20 differentially expressed genes within Melanoma-c3 cluster. (*D*) Results of the GO-BP enrichment analysis for the top 20 differentially expressed genes within Melanoma-c4 cluster. (*E*) GSEA plot of the enrichment of the Hallmark Epithelial-Mesenchymal Transition (EMT) pathway between cancer-c4 cluster and cancer-c3 cluster. (*F*) Ligand-receptor cellular interactions between CTLs (CD8T-cx) and cancer cells (Cancer-c4 or -c3) in the coculture of OVA cancer cells and CTLs. Only the interactions with a p value less than 0.01 were displayed. The color represents the communication probability as indicated by the color bar.

**Movie S1 (separate file).** Representative movie of the co-culture between CTLs and OVA cancer cells.

**Movie S2 (separate file).** Representative movie of the co-culture between CTLs and WT cancer cells.

**Movie S3 (separate file).** Representative movie of collagen fiber in the co-culture of CTLs and OVA cancer cells at 0 h.

**Movie S4 (separate file).** Representative movie of collagen fiber in the co-culture of CTLs and WT cancer cells at 0 h.

**Movie S5 (separate file).** Representative movie of collagen fiber in the co-culture of CTLs and OVA cancer cells at 6 h.

**Movie S6 (separate file).** Representative movie of collagen fiber in the co-culture of CTLs and WT cancer cells at 6 h.

**Movie S7 (separate file).** Simulated T cell motility in a simulation of multiple cancer clusters.

**Movie S8 (separate file).** Simulated T cell motility in a simulation of a single cancer cluster.

**Dataset S1 (separate file).** Statistical source data.

**Dataset S2 (separate file).** Statistical source data.

**Dataset S3 (separate file).** Statistical source data.

**Dataset S4 (separate file).** Statistical source data.

**Dataset S5 (separate file).** Statistical source data.
